# Cell growth rate dictates the onset of glass to fluid-like transition and long time super-diffusion in an evolving cell colony

**DOI:** 10.1101/174599

**Authors:** Abdul N Malmi-Kakkada, Xin Li, Himadri S. Samanta, Sumit Sinha, D. Thirumalai

## Abstract

Collective migration dominates many phenomena, from cell movement in living systems to abiotic self-propelling particles. Focusing on the early stages of tumor evolution, we enunciate the principles involved in cell dynamics and highlight their implications in understanding similar behavior in seemingly unrelated soft glassy materials and possibly chemokine-induced migration of CD8^+^ T cells. We performed simulations of tumor invasion using a minimal three dimensional model, accounting for cell elasticity and adhesive cell-cell interactions as well as cell birth and death to establish that cell growth rate-dependent tumor expansion results in the emergence of distinct topological niches. Cells at the periphery move with higher velocity perpendicular to the tumor boundary, while motion of interior cells is slower and isotropic. The mean square displacement, Δ(*t*), of cells exhibits glassy behavior at times comparable to the cell cycle time, while exhibiting super-diffusive behavior, Δ(*t*) ≈ *t^α^* (*α* > 1), at longer times. We derive the value of *α* ≈ 1.33 using a field theoretic approach based on stochastic quantization. In the process we establish the universality of super-diffusion in a class of seemingly unrelated non-equilibrium systems. Super diffusion at long times arises only if there is an imbalance between cell birth and death rates. Our findings for the collective migration, which also suggests that tumor evolution occurs in a polarized manner, are in quantitative agreement with *in vitro* experiments. Although set in the context of tumor invasion the findings should also hold in describing collective motion in growing cells and in active systems where creation and annihilation of particles play a role.

## I. INTRODUCTION

The strict control of cell division and apoptosis is critical for tissue development and maintenance [1]. Dysfunctional cell birth and death control mechanisms lead to several physiological diseases including cancers [2]. Together with genetic cues controlling birth-death processes, mechanical behavior of a collection of cells is thought to be of fundamental importance in biological processes such as embryogenesis, wound healing, stem cell dynamics, morphogenesis, tumorigenesis and metastasis [3–8]. Due to the interplay between birth-death processes and cell-cell interactions, we expect that collective motion of cells ought to exhibit unusual non-equilibrium dynamics, whose understanding might hold the key to describing tumor invasion and related phenomena. Interestingly, characteristics of glass-like behavior such as diminished motion (jamming) of a given cell in a dense environment created by neighboring cells (caging effect), dynamic heterogeneity, and possible viscoelastic response have been reported in confluent tissues [9, 10]. Using imaging techniques that track cell motions, it has been shown that in both two (kidney cells on a flat thick polyacrylamide gel [9, 11]) and three dimensions (explants from zebrafish embedded in agarose [12]) the mean displacement exhibits sub-diffusive behavior, reminiscent of dynamics in supercooled liquids at intermediate time scales. This behavior, which can be rationalized by noting that the core of a growing collection of cells is likely to be in a jammed state, is expected on time scales less than the cell division time.

A theory to capture the essence of tumor invasion must consider the interplay of the cell mechanics, adhesive interaction between cells, and the dynamics associated with cell division and apoptosis, over a wide range of time scales. In an attempt to capture collective dynamics in cells a number of models based on cellular automaton [13, 14], vertex and Voronoi models [15–19], subcellular element model [20], cell dynamics based on Potts model [21, 22], and phase field description for collective migration [23, 24] have been proposed. Previous works have investigated a number of two-dimensional (2D) models in various contexts [9, 16, 25] including probing the dynamics in a homeostatic state where cell birth-death processes are balanced [26, 27]. Existing threedimensional (3D) models focus solely on tumor growth kinetics, spatial growth patterns [28, 29] or on cell migration at low cellular density on time scales shorter than the cell division time [30-32]. A recent interesting study [26] shows that cell dynamics in a confluent tissue is always fluidized by cell birth and death processes, on time scales comparable to cell division time. A more recent two-dimensional model [27] investigates glass-to-liquid transition in confluent tissues using simulations. However, both these instructive models [26, 27] focus on the steady state regime where the number of tumor cells is kept constant by balancing the birth and death rates. Consequently, they do not address the non-equilibrium dynamics of the evolving tumor in the early stages, which is of great interest in cancer biology [33–35].

Here, we use a minimal physical 3D model that combines both cell mechanical characteristics, cell-cell adhesive interactions and variations in cell birth rates to probe the non-steady state tumor evolution. Such a model, which has the distinct advantage that it can be generalized to include mutational effects naturally, was first introduced by Drasdo and Hohme [28]. One of our primary goals is to understand quantitatively the complex invasion dynamics of tumor into a collagen matrix, and provide a mechanism for the observation of super-diffusive behavior. We use free boundary conditions for tumor evolution to study dynamical fingerprints of invasion in order to quantitatively compare the results to experimental observations. We model the proliferation behavior of tumor cells, and investigate the effect of pressure dependent growth inhibition. Good agreement between our results and *in vitro* experiments on three-dimensional growth of multicellular tumor spheroids lends credence to the model. On time scales less than the cell division time, the dynamics of cell movement within the tumor exhibits glassy behavior, reflected in a sub-diffusive behavior of the mean square displacement, Δ(*t*). However, at times exceeding cell division time we find super-diffusive behavior with Δ(*t*) ~ *t^α^* (*α* = 1.26 ± 0.05). The duration for which sub-diffusion persists decreases as the cell growth rate increases, in sharp contrast to the dynamics in confluent tissues. Detailed analyses of the individual cell trajectories reveal complex heterogeneous spatial and time dependent cell migration patterns, thus providing insights into how cells are poised for invasion into regions surrounding the tumor. We find that activity due to cell division coupled with cell mechanical interactions plays a critical role in the non-equilibrium dynamics and the physical structure of the polarized tumor invasion process. The dynamical properties of cells in our model share considerable similarities to those found in non-living soft materials such as soap foams and toothpaste [36]. In all these cases the transition from a glass-like behavior to super diffusion occurs as a result of cell growth and death (or creation and destruction of particles), resulting in non-conservation of number density without the possibility of reaching homeostasis. In other words, the non-equilibrium dynamics arising due to forces that result from key biological events (cell birth and death) that we have investigated here are qualitatively different from dynamics in systems which do not take into account such forces.

## II. MULTICELLULAR TUMOR GROWTH MODEL

We simulated the spatiotemporal dynamics of a multicellular tumor using a three dimensional (3D) agent-based model in which the cells in the tumor are represented as interacting objects. In this model, the cells grow stochastically as a function of time and divide upon reaching a critical size. The cell-to-cell interaction is characterized by direct elastic and adhesive forces. We also consider cell-to-cell and cell-to-matrix damping as a way of accounting for the effects of friction experienced by a moving cell due to other cells, and by the extracellular matrix (ECM) (or collagen matrix), respectively.

Each cell is modeled as a deformable sphere with a time dependent radius. Several physical properties such as the radius, elastic modulus, membrane receptor and ligand concentration, adhesive interaction, characterize each cell. Following previous studies [28, 29, 37], we use the Hertzian contact mechanics to model the magnitude of the elastic force between two spheres of radii *R*_*i*_ and *R_j_* (Fig. 1a),

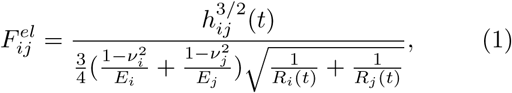

where *E_i_* and *ν_i_*, respectively, are the elastic modulus and Poisson ratio of cell *i*. The overlap between the spheres, if they interpenetrate without deformation, is *h_ij_*, which is defined as max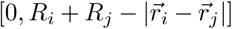 with 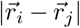 being the center-to-center distance between the two spheres (see Fig. 1a). The repulsive force in Eq. (1) is valid for small virtual overlaps such that *h_ij_* ≪ min[*R_i_*, *R_j_*], and is likely to underestimate the actual repulsion between the cells [29]. Nevertheless, the model incorporates measurable mechanical properties of the cell, such as *E_i_* and *v_i_*, and hence we use this form for the repulsive force.

**FIG. 1.**
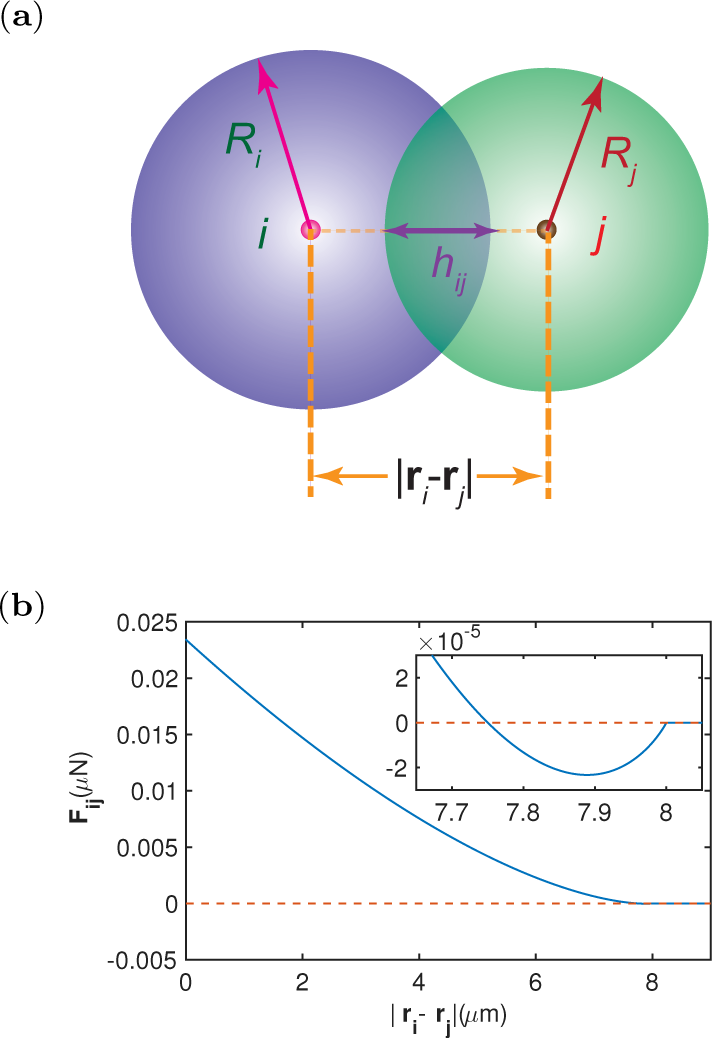
(a) Illustration of two interpenetrating cells *i* and *j* with radii *R_i_* and *R_j_*, respectively. The distance between the centers of the two cells is |**r**_*i*_ – **r**_*j*_|, and their overlap is *h_ij_*. (b) Force on cell *i* due to *j*, **F**_*ij*_, for *R*_*i*_ = *R_j_* =4 *μm* using mean values of elastic modulus, poisson ratio, receptor and ligand concentration (see Table I). F*ij* is plotted as a function of distance between the centers of the two cells. Inset shows the region where **F**_*ij*_ is attractive. When |**r**_*i*_ – **r**_*j*_ ≥ *R_i_* + *R_j_* = 8 *μm* the cells are no longer in contact, and hence, **F**_*ij*_ = 0.

**Table 1:**
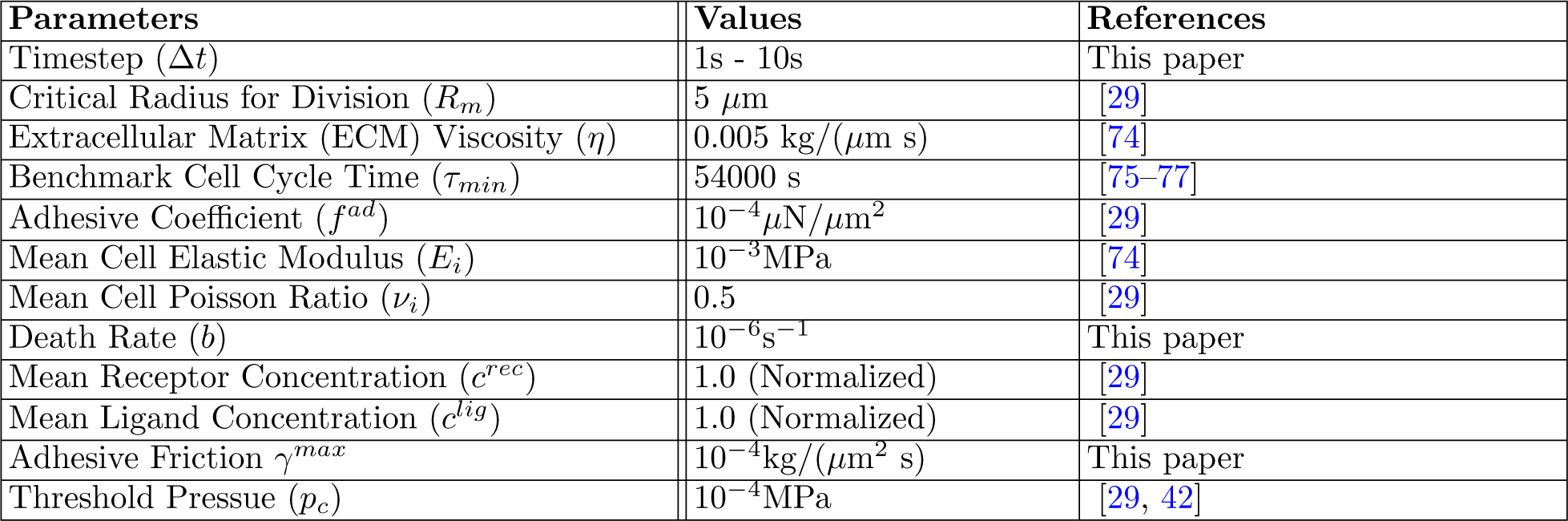
The parameters used in the simulation.

| Parameters | Values | References |
| --- | --- | --- |
| Timestep ($\Delta t$) | 1s - 10s | This paper |
| Critical Radius for Division ($R_m$) | 5 $\mu\text{m}$ | [29] |
| Extracellular Matrix (ECM) Viscosity ($\eta$) | 0.005 kg/($\mu\text{m s}$) | [74] |
| Benchmark Cell Cycle Time ($\tau_{min}$) | 54000 s | [75–77] |
| Adhesive Coefficient ($f^{ad}$) | $10^{-4} \mu\text{N}/\mu\text{m}^2$ | [29] |
| Mean Cell Elastic Modulus ($E_i$) | $10^{-3} \text{MPa}$ | [74] |
| Mean Cell Poisson Ratio ($\nu_i$) | 0.5 | [29] |
| Death Rate ($b$) | $10^{-6} \text{s}^{-1}$ | This paper |
| Mean Receptor Concentration ($c^{rec}$) | 1.0 (Normalized) | [29] |
| Mean Ligand Concentration ($c^{lig}$) | 1.0 (Normalized) | [29] |
| Adhesive Friction $\gamma^{max}$ | $10^{-4} \text{kg}/(\mu\text{m}^2 \text{s})$ | This paper |
| Threshold Pressue ($p_c$) | $10^{-4} \text{MPa}$ | [29, 42] |

Cell adhesion, mediated by receptors on the cell membrane, is the process by which cells interact and attach to one another. For simplicity, we assume that the receptor and ligand molecules are evenly distributed on the cell surface. Consequently, the magnitude of the adhesive force, 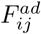, between two cells i and j is expected to scale as a function of their contact area, *A_ij_* [38]. We estimate 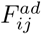 using [29],

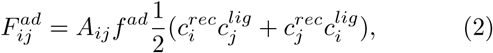

where the 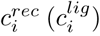 is the receptor (ligand) concentration (assumed to be normalized with respect to the maximum receptor or ligand concentration so that 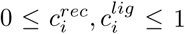. The coupling constant f^*ad*^ allows us to rescale the adhesion force to account for the variabilities in the maximum densities of the receptor and ligand concentrations. We calculate the contact surface area, *A_ij_*, using the Hertz model prediction, *A_ij_ =πh_ij_R_i_R_j_*/(*R_i_* + *R_j_*). The Hertz contact surface area is smaller than the proper spherical contact surface area. However, in dense tumors many spheres overlap, and thus the underestimation of the cell surface overlap may be advantageous in order to obtain a realistic value of the adhesion forces [29].

Repulsive and adhesive forces considered in Eqs.(1) and (2) act along the unit vector 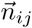 pointing from the center of cell *j* to the center of cell *i* (Fig. 1a). The force exerted by cell *i* on cell *j*, **F**_*ij*_, is shown in Fig. 1b. The total force on the *i^th^* cell is given by the sum over its nearest neighbors (*NN*(*i*)),

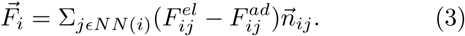

We developed a distance sorting algorithm to efficiently provide a list of nearest neighbors in contact with the *i^th^* cell for use in the simulations. For any given cell, *i*, an array containing the distances from cell *i* to all the other cells is initially created. We then calculated,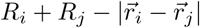 and sorted the cells *j* satisfying 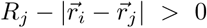 - a necessary condition for any cell *j* to be in contact with cell *i*.

## III. SIMULATIONS

### Equations of Motion

The spatial dynamics of the cell is computed based on the equation of motion [29, 39, 40] for a cell of mass *m_i_*,

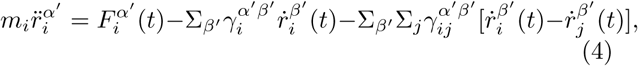

where the Greek indices [*α′,β′*] = [*x,y, z*] are for coordinates, and the Latin indices [*i, j*] = [1, 2, …, *N*] are the cell indices. In Eq. (4), 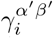 is the cell-to-medium friction coefficient, and 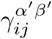 is the cell-to-medium friction coefficient The adhesive and repulsive forces are included in the term 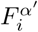. The cell-to-ECM friction coefficient is assumed to be given by the Stokes relation,

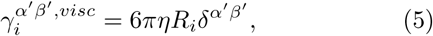

based on the friction of a sphere in a medium of viscosity *η*. Here, 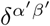 is the Kronecker delta.

Because the Reynolds number for cells in a tissue is small [39], overdamped approximation is appropriate implying that the neglect of the inertial term 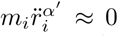is justified [29] (see Appendix A for further discussion). Since additional adhesive forces are also present, cell movement is further damped [41]. We simplify the equations of motion (Eq. (4)), by replacing the intercellular drag term with a modified friction term, given that the movement of the bound cells is restricted. The modified friction term will contribute to the diagonal part of the damping matrix with 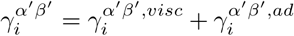 where,

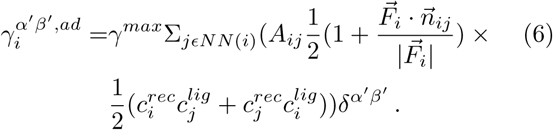

Notice that the added friction coefficient 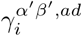 is proportional to the cell-to-cell contact surface, implying that a cell in contact with multiple cells would move less. The non-isotropic nature of the adhesive friction is evident from the factor 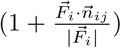 where the maximum contribution occurs when the net force 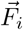 is parallel to a given unit vector, 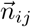, among the nearest neighbors. With these approximations, the equations of motion (Eq. (4)) are now diagonal,

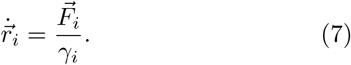

### Cell Cycle

In our model, cells can be either in the dormant (*D*) or in the growth (*G*) phase. We track the sum of the normal pressure that a particular cell *i* experiences due to contact with its neighbors, using,

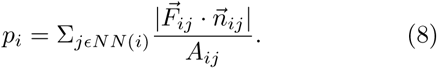

If the local pressure, *p_i_*, exceeds a critical limit (*p_c_*) the cell stops growing and enters the dormant phase (see the left panel in Fig. 2). For growing cells, their volume increases at a constant rate r_V_. The cell radius is updated from a Gaussian distribution with the mean rate 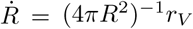. Over the cell cycle time *τ*,

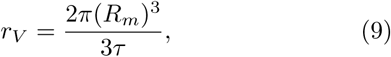

where *R_m_* is the mitotic radius. A cell divides once it grows to the fixed mitotic radius. To ensure volume conservation, upon cell division, we use *R_d_* = *R_m_*2^−1/3^ as the radius of the daughter cells (see the right panel in Fig. 2). The two resulting cells are placed at a center-to-center distance *d = 2R_m_*(1 – 2^−1/3^). The direction of the new cell location is chosen randomly from a uniform distribution on the unit sphere. One source of stochastic-ity in the cell movement in our model is due to random choice for the mitotic direction. Together with stochas-ticity in the cell cycle duration, we obtain fairly isotropic tumor spheroids.

**FIG. 2.**
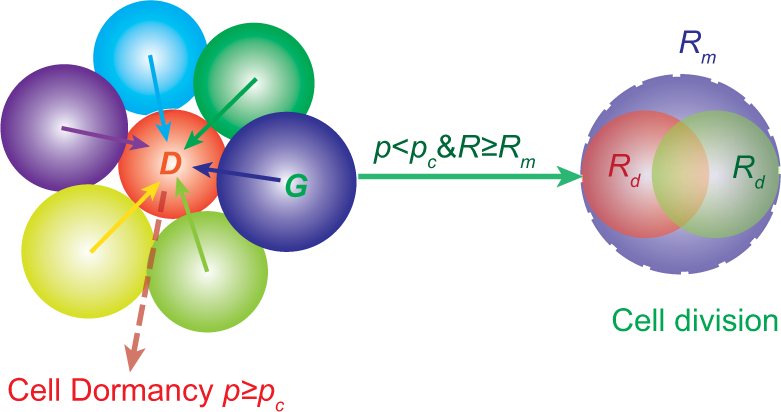
Cell dormancy (left panel) and cell division (right panel). If the local pressure *p_i_* that the *i*^th^ cell experiences (due to contacts with the neighboring cells) exceeds the critical pressure *p_c_*, it enters the dormant state (*D*). Otherwise, the cells grow (*G*) until they reach the mitotic radius, *R_m_*. At that stage, the mother cell divides into two identical daughter cells with the same radius *R_d_*. We assume that the total volume upon cell division is conserved. A cell that is dormant at a given time can transit from that state at subsequent times.

### Tumor invasion distance

The invasion or spreading distance of the growing tumor, Δr(*t*), is determined by measuring the average distance from the tumor center of mass 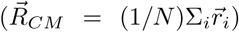 to the cells at the tumor periphery,

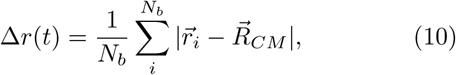

where the sum *i* is over *N_b_*, the number of cells at the tumor periphery. In order to find the cells at the tumor periphery (*N_b_*), we denote the collection of all cells *N* as a set of vertices {1, 2, 3,………*N*} in 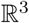 where the vertices represent the center point of each cell. We generate a 3D structure of tetrahedrons using these vertices, where each tetrahedron is comprised of 4 vertices. Let *T* be the total number of tetrahedrons. Since each tetrahedron has 4 faces, the total number of faces is 4*T*. Any face that is not on the boundary of the 3D structure is shared by 2 tetrahedrons but the boundary faces are not shared. Thus, our aim is to find the set of unshared boundary faces out of the total number, 4*T*, of faces. Once the boundary faces are obtained, we know the list of vertices, and hence the positions of cells at the tumor periphery, allowing us to compute Δ*r*(*t*)in Eq. (10).

## IV. RESULTS

### Calibration of the model parameters

We compare the normalized volume of the growing tumor to experimental data [42], as a way of assessing if the parameters (Table I) used in our model are reasonable. The tumor volume, *V*(*t*), normalized by the initial volume of the spheroid (*V*_0_), was tracked experimentally using colon carcinoma cells [42] through experimental methods that are very different from the way we simulated tumor growth. The tumor growth was measured by imposing stress [42], known to inhibit cancer growth [43]. These effects are included in our model, which allows us to make quantitative comparisons between our simulations and experiments. The tumor volume is obtained in the simulation from, *R_g_*, the radius of gyration,

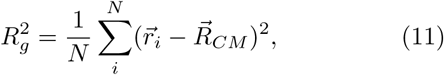

where N is the total number of cells. The volume *V*(*t*) is given by 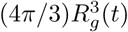. Our simulation of the growth of the spheroid tumor volume in the early stages is in good agreement with experimental data (see Fig. 3a). Thus, our model captures quantitative aspects of tumor growth.

**FIG. 3.**
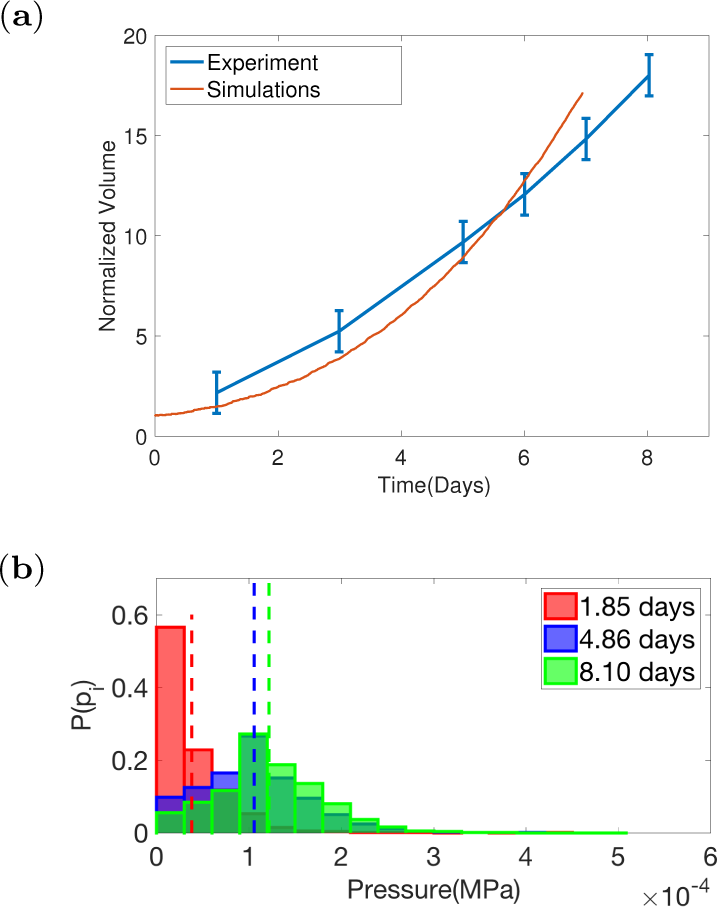
Normalized volume, *V(t)/V*_0_, of a tumor spheroid as a function of time. The result of simulation (red) agrees nearly quantitatively with experimental data obtained for the tumor spheroid growth at an applied pressure of 500Pa [42]. We used a critical pressure *p_c_* = 100Pa and cell division time of *τ* = *τ_min_* in these simulations. The reason for comparing the results from the *pc* = 100Pa simulations with the growth dynamics obtained in the colon carcinoma cells with an external pressure of 500Pa is explained in the text. (b) Distribution of pressure as a function of total growth time with cell division time *τ* = *τ_min_* = 0.625 days. The mean values are indicated by the dashed lines.

In the experiment [42], *V(t)/V*_0_ was measured for external pressure ranging from 0 – 20kPa. In Fig. (3a), we compared our simulation results with the 500Pa result from experiments [42]. This is rationalized as follows. Unlike in experiments, the pressure is internally generated as the tumor grows (Fig. 3b) with a distribution that changes with time. The mean value of the pressure (see dashed lines in Fig. 3b) at the longest time is ≈ 100Pa. Thus, it is most appropriate to compare our results obtained using *p_c_* = 100Pa with experiments in which external pressure is set at 500Pa.

### Predicted pressure-dependent growth dynamics is consistent with experiments

Visual representation of the tumor growth process generated in simulations is vividly illustrated in the movie (see Movies 1 and 2) in the Appendix B. Snapshots of the evolving collection of cells at different times are presented in Figs. 4a-d. As the tumor evolves, the cells aggregate into a spheroidal shape due to cell division plane being isotropically distributed (Fig. 4d and the movie in the Appendix B). In spheroidal cell aggregates, it is known that pressure inhibits cell proliferation [42, 44, 45]. We expect the pressure (see Eq. 8) experienced by the cells in the interior of the tumor to be elevated due to crowding effects, causing the cells to enter a dormant state if the pressure from the neighbors reaches a preset threshold value, *p_c_*. Tumor growth behavior is strongly dependent on the value of *p_c_* (see Fig. 4e). At *p_c_* = 10^−3^ MPa, the total number (*N*(*t*)) of tumor cells during growth is well approximated as an exponential *N*(*t*) ∝ *exp*(*const × t*). As *p*c** is lowered, implying growth is inhibited at smaller pressures, increase in the tumor size is described by a power law, *N*(*t*) ∝ *t^β^*, at long timescales, while *N*(*t*) retains exponential growth at early stages (see the inset of Fig. 4e). Our simulations also show that *β* is *p_c_*- dependent, increasing as *p*c** increases. Power-law growth in 3D tumor spheroid size has been observed in many tumor cell lines with *β* varying from one to three [46–50]. The overall growth of the tumor slows down as the value of pressure experienced by cells increases, which is also consistent with recent experimental results [45]. The known results in Fig. 4e, are in near quantitative agreement with several experiments, thus validating the model.

**FIG. 4.**
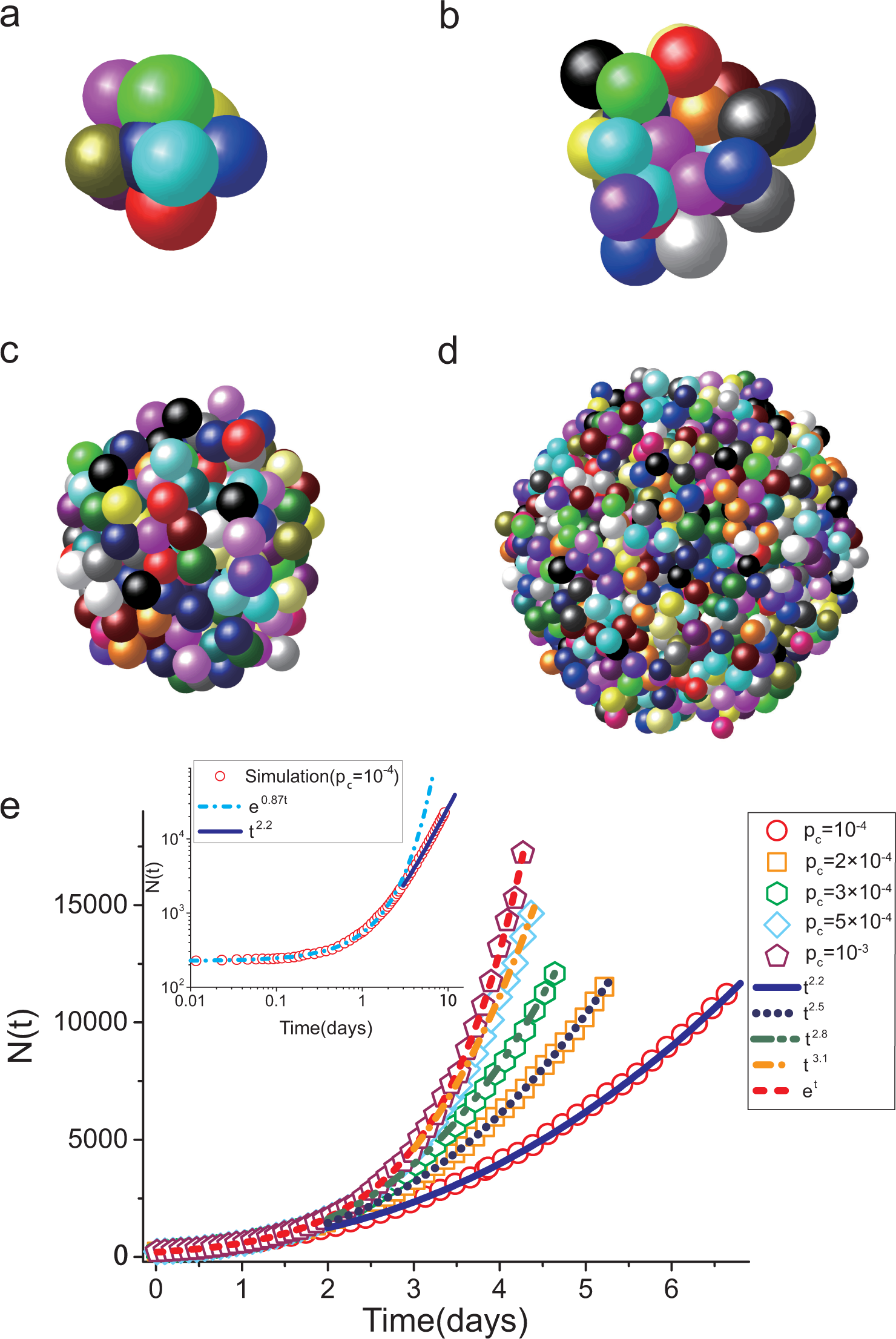
Dynamics of Tumor growth. **(a-d)** show instantaneous snapshots of the tumor during growth at different times. Each cell is represented by a sphere. There are approximately 2,000 cells in **(d)**. The color of each cell is to aid visualization. In figures **(a-d)** the cell sizes are rescaled for illustration purposes only. Note that even at *t* = 0 **(a)** the sizes of the cells are different because they are drawn from a Gaussian distribution. **(e)**, The total number of tumor cells, *N*(*t*), as a function of time at different values of the threshold pressure *p*_c_, which increases from bottom to top (10^−4^, 2 × 10^−4^, 3 × 10^−4^, 5 × 10^−4^, 10^−3^ MPa). The dashed red line is an exponential function while other lines show power-law behavior *N*(*t*) ≈ *t^β^* where *β* ranges from 2.2, 2.5, 2.8 to 3.1 (from bottom to top). The inset in **(e)** shows *N*(*t*) with *pc* = 10^−4^ on a log-log scale with both exponential and power law fits. The dash-dot curve in the inset is an exponential function while the power-law trend is illustrated by the solid line. The average cell cycle time *τ = τ_min_* = 54,000s and other parameter values are taken from Table I.

### Cell motility within the tumor spheroid

Using direct imaging techniques it has become possible to monitor the overall invasion of the tumor as well as the movement of individual cells within the spheroid [51, 52]. In order to compare our results to experiments we calculated the mean square displacement, 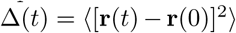, of individual cells. By tracking the movement of all the initial cells within the tumor, we calculated Δ(*t*) by averaging over hundreds of trajectories. The growth rate-dependent Δ(*t*) (displayed in Fig. 5a on a log-log scale) shows that there is a rapid increase in Δ(*t*) at early times (*t* ≤ 0.01 *τ_min_*, where *τ_min_* is the benchmark value of the cell cycle time given in Table I) because the cells move unencumbered, driven by repulsive interaction with other cells. At intermediate timescales (0.01*τ_min_* < *t* < *τ* with *τ* being the average cell cycle time), Δ(*t*) exhibits sub-diffusive behavior (Δ(*t*) ~ t^s^ with s < 1). The signatures of the plateaus in Δ(*t*) (together with other characteristics discussed later) in this time regime indicate that cells are caged by the neighbors (see the left inset in fig. 5a), and consequently undergo only small displacements. Such a behavior is reminiscent of a supercooled liquid undergoing a glass transition, as illustrated in colloidal particles using direct imaging as their densities approach the glass transition [53, 54]. As *τ* increases, the plateau persists for longer times because of a decrease in the outward stress, which slows the growth of the tumor. When *t* exceeds *τ* the average cell doubling time, the Δ(*t*) exhibits super-diffusive motion Δ(*t*) ≈ *t^α^* (*α* > 1). In order to determine *α* we performed multiple simulations and calculated Δ(*t*) for each of them by generating a large number of trajectories. All the independent simulations show that Δ(*t*) has the characteristic plateau at intermediate times followed by super-diffusion at long times in Fig. 5b. From each of these simulations we determine *α*, whose distribution is shown in Fig. 5c. In all cases, we find that *α* is greater than unity. The estimate from the distribution in Fig. 5c is *α* = 1.26 ± 0.05 where 0.05 is the standard deviation.

**FIG. 5.**
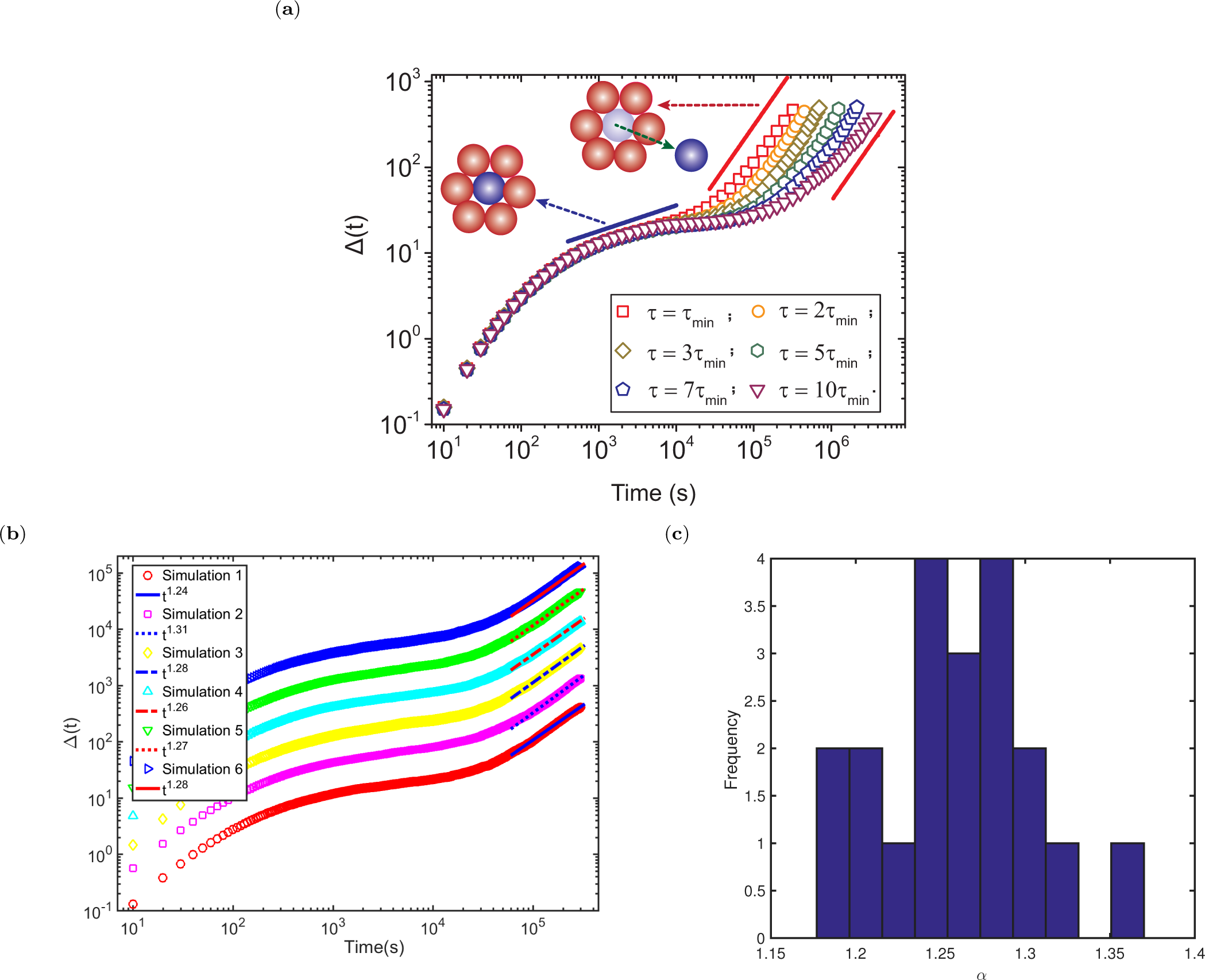
Super-diffusive behavior in Δ(*t*) at long times. (**a**) The mean-squared displacement (Δ(*t*)) of cells. From top to bottom, the curves correspond to increasing average cell cycle time (*τ* is varied from *τ_min_* to 10*τ_min_* where *τ_min_* = 54, 000 seconds). Time taken for reaching the super-diffusive regime increases by increasing τ. The blue and red lines have a slopes of 0.33, and 1.3, respectively. The sub-diffusive (super-diffusive) behavior corresponds to dynamics in the intermediate (long) times. The left and middle inset illustrate the “cage effect” and “cage-jump” motion, respectively. The unit for y-axis is (*μm^2^*). (**b**) Fit of the mean-square displacement, Δ(*t*) to *t^α^* for several simulation runs. Each Δ(*t*) plot is averaged over ≈ 300 cell trajectories. The six Δ(*t*) plots consistently show *α >* 1.20 over ≈ 1800 cell trajectories. The plots are separated for clarity. (**c**) Distribution of a values from multiple independent simulations. Mean value of a is 1.26 with a standard deviation of ±0.05.

On long time scales, cells can escape the cage created by their neighbors, as illustrated in the middle inset of Fig. 5a. Our observation of super-diffusion in Δ(*t*) at long times agrees well with the experimental result (*α* ≈ 1.40 ± 0.04) obtained for fibrosarcoma cells in a growing tumor spheroid [33]. The onset of super-diffusive behavior in Δ(*t*) shifts to earlier times as we decrease the average cell cycle time (see Fig. 5a), implying that cell division is the mechanism resulting in super-diffusion (see below for further discussion of this crucial finding).

We provide another rationale for robustness of the long time super diffusive behavior. This comes from examining the time-dependent changes in the invasion distance, Δ*r*(*t*) in Eq.10. The finding that the invasion distance does not increase as a function of time with exponent Δ*r ∝ t*^0.5^ but rather at a higher exponent at the long time regime necessarily implies that cells do not execute random walk motion (see Ref. [33]). The dependence of Δ*r*(*t*) on t in Fig. 6 shows that for *t* ≤ *τ_min_*, the invasion radius is roughly constant. As cells divide the tumor invasion distance, Δr(*t*), increases as *t^ξ^* with *ξ* ~ 0.63 (implies *α* ≈ 1.26) for *t* > *τ_min_*, a value that is not inconsistent with experiments [33].

**FIG. 6.**
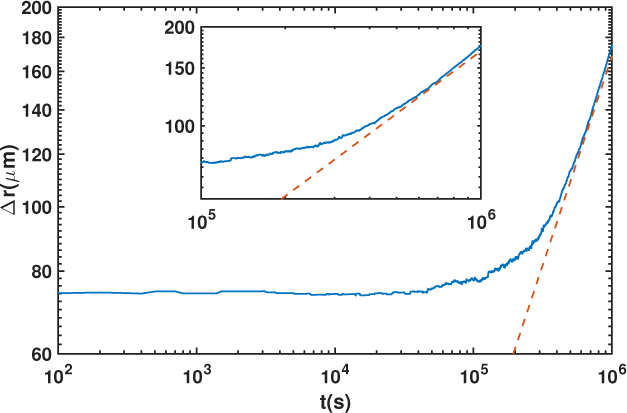
The invasion distance, Δ*r*(*t*) (Eq. 10), as a function of time. The exponent of the invasion distance, as indicated by the dashed line, determined using Δ*r* ∝ *t^ξ^* is *ξ* ≈ 0.63. Note that this value and the one extracted from experiments [331 are in reasonable agreement. The inset shows Δ*r*(*t*) for *t* > *τ*_*min*_ for average cell cycle time *τ* = 1*τ_min_*.

### Theoretical predictions

In order to understand the role of cell growth and apoptosis in the observed sluggish dynamics at intermediate times and super-diffusive behavior at long times, we developed a theory to study the dynamics of a colony of cells in a dissipative environment (Appendix C). The interactions between cells contain both attractive (adhesive) and excluded volume terms. Starting from the Langevin equation describing the dynamics of the *i^th^* cell, and incorporating the birth reaction, 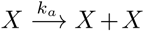*X* with the rate constant *k_a_* (= 1/*τ*) for each cell, and the apoptosis reaction 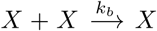 with the rate *k_b_*, an equation for the time dependence of the density *ρ*(**k**, *t*) (Eq. C3 in the Appendix C) can be derived. The cell division and apoptosis processes drive the system far from equilibrium, thus violating the Fluctuation Dissipation Theorem (FDT). As a consequence, we cannot use standard methods used to calculate response and correlation functions from which the t-dependence of Δ(*t*) can be deduced. To overcome this difficulty we used the Parisi-Wu stochastic quantization method [55] in which the evolution of *ρ*(**k**, *ω*) (*ω* is the frequency) is described in a fictitious time in which FDT is preserved.

From the analysis of the resulting equation (Appendix C contains the sketch of the calculations) the scaling of Δ(*t*) may be obtained as,

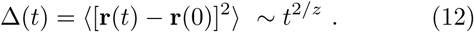

In the intermediate time regime, *z* = 5/2, implying Δ(*t*) ≈ *t*^4/5^. The predicted sub-diffusive behavior of Δ(*t*) is qualitatively consistent with simulation results. It is likely that the differences in the scaling exponent between simulations (2/*z* ≈ 0.33) and theoretical predictions (2/*z* ≈ 0.80) in this non-universal time regime may be due to the differences in the cell-to-cell interactions used in the two models.

In the long time limit, the cell birth-death process (the fourth term in Eq. C3) dominates the interactions between cells. As a result, we expect that the exponent *2/*z** should be universal, independent of the forces governing cell motility. Our theory predicts that *z* = 3/2, which shows that Δ(*t*) ≈ *t*^4/3^, in excellent agreement with the simulations (Fig. 5a) and experiments [33]. It is clear from our theory that the interaction-independent biologically important birth-death processes drive the *observed fluidization* during tumor (or tissue) development, resulting in super-diffusive cell motion at long times. The underlying mechanism for obtaining super-diffusive behavior is that cells must move persistently in a given direction for a long time leading to polarized tumor growth, ultimately resulting in invasion driven predominantly by birth. We provide additional numerical evidence for this assertion below.

### Dependence of relaxation times on cell cycle time

We first characterized the structural evolution as the tumor evolves. In order to assess the spatial variations in the positions of the cells as the tumor grows, we calculated the pair correlation function using,

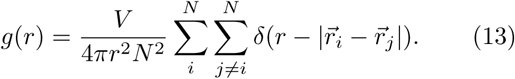

The pair correlation functions (Fig. 7), at different cell cycle times (*τ*), show that at longer cell cycle times, the cells are packed more closely. There is a transition from a liquid-like to a glass-like structure as *τ* is increased, as indicated by the peaks in *g*(*r*).

**FIG. 7.**
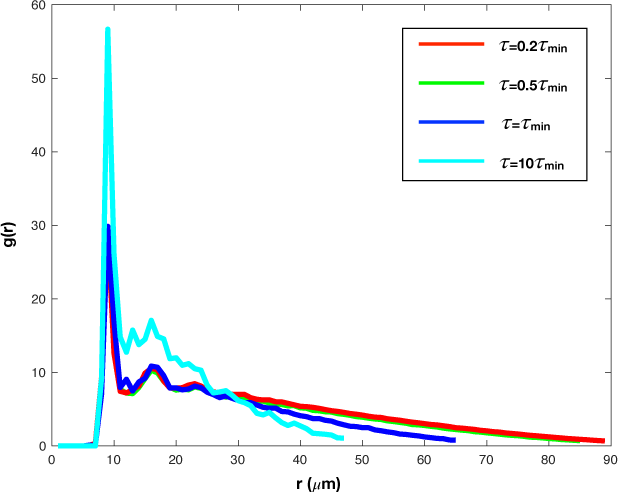
Pair correlation function at four different cell division times (*τ*): 0.2*τ_min_* (red), 0.5*τ_min_* (green), *τ_min_* (blue), and 10r*τ_min_* (cyan). The cells are packed more closely at longer cell cycle times as reflected by the sharper peak for the cyan line compared to the others. The distance, *r*, at which *g(r)* approaches zero is considerably smaller for *τ*= 10*τ_min_*. The distance at which the first peak appears is ≈ *2R_m_* ≈ 10*μm* (Table I), which implies that despite being soft the cells in the interior are densely packed, as in a body centered cubic lattice.

To further quantify the fluidization transition driven by cell birth-death processes, we calculated the isotropic self-intermediate scattering function 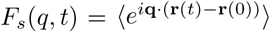 at |**q**| = 2π/*r*_0_, where *r*_0_ is the position of the first maximum in the pair correlation function (see Fig. 7). The average is taken over all the initial cells, which are alive during the entire simulation time and the angles of **q**. We note that *F_s_*(*q*,*t*) exhibits a two-step relaxation process (Fig. 8a) characterized by two time scales. The initial relaxation time, corresponding to the motion of cells in a cage formed by neighboring cells, depends only weakly on the cell cycle time. The second relaxation time (*τ_α_*), extracted by fitting *F_s_*(*q*,*t*) to an exponential function 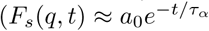, see colored solid lines in Fig. 8a), depends strongly on the average cell cycle time. As in the relaxation of supercooled liquids, **τ_α_** is associated with the collective motion of cells leaving the cage [56, 57]. As the average cell cycle time is reduced, *T_a_* decreases (see Fig. 8b), and the *F_s_*(*q*,*t*) begins to approach a single relaxation regime, as expected for a normal fluid. The second relaxation process in *F_s_*(*q*,*t*) (Fig. 8a) can be collapsed onto one master curve by rescaling time by **τ_α_*,* resulting in the independence of *F_s_ (q, t)* on the cell cycle time (Fig. 8c). We surmise that the cage relaxation is driven by the same mechanism (the cell birth-death processes) that gives rise to super-diffusive behavior in Δ(*t*).

**FIG. 8.**
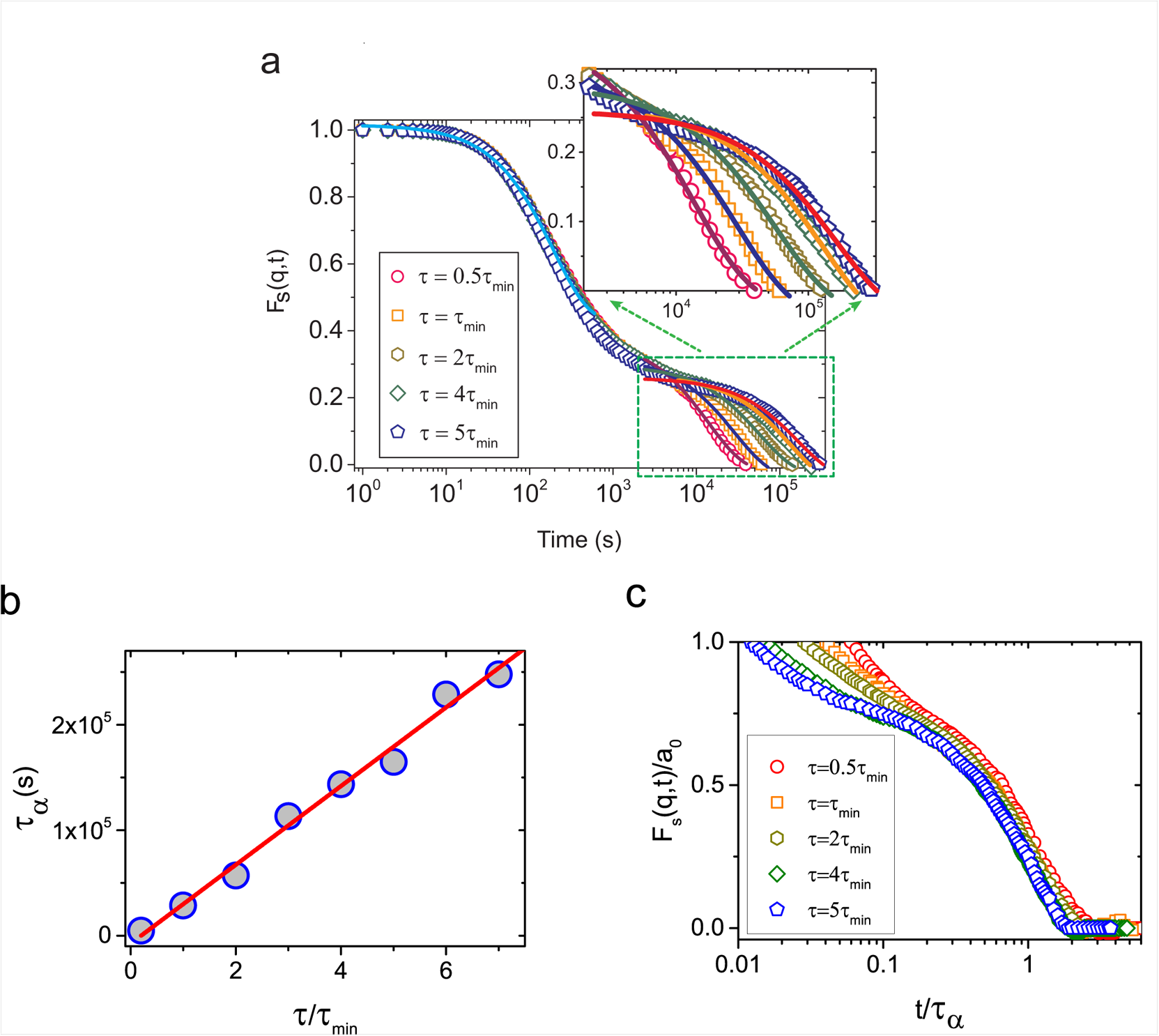
Self intermediate scattering function at different cell cycle times. (**a**) The self-intermediate scattering function, *F_s_*(*q,t*), shows that relaxation occurs in two steps. From left to right, the second relaxation for *F_s_*(*q*,*t*) slows down as *τ* increases (from 0.5*τ_min_* to 5*τ_min_*). The solid lines are exponential fits. The upper inset shows a zoom-in of the dashed-line rectangle at long timescales. (**b**) The second relaxation time *τ_α_* obtained from (**a**) as a function of cell division time (rescaled by *τ_min_*). The red solid line is a linear fit *(*τ_α_** ∝ 0.69*τ*). (**c**) The rescaled self-intermediate scattering function *F_s_*(*q,t*)/*a*_0_ as a function of the rescaled time *t*/*τ_α_*.

### Diffusion of tracer cells

Elsewhere [27] using a twodimensional model Δ(*t*) was computed for cells as well as tracer cells. In their study, using periodic boundary conditions, the choice of birth and death rates is such that in the long time limit homeostasis is always reached where birth and death of cells is balanced. It was found that Δ( *t*) for live cells (those that can be born and die) show a plateau at intermediate times followed by normal diffusion (Δ(*t*) ~ *t*) at long times. In contrast, Δ_*tr*_(*t*) the mean squared displacement computed for tracer cells (ones which have all the characteristics of cells except they are alive throughout the simulations and do not grow or divide) shows no caging effects but grows linearly with time [27], suggestive of normal diffusion. In light of this dramatically different behavior reported in [27], we performed simulations using our model by including 100 randomly placed tracer cells. The interactions between the tracer cells with each other and the cells that undergo birth and death are identical. The calculated dependence of Δ_*tr*_(*t*) for tracer cells, as a function of *t* (Fig. 9a) is qualitatively similar to that for cells (compare Fig. 5a and Fig. 9a). In particular at varying values of *τ*, Δ*_tr_*(*t*) exhibits a plateau followed by super-diffusive behavior, 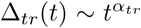 at long times. However, we find that *α_tr_* (> 1.4) depends on *τ* in contrast to the universal exponent for cell dynamics. Similarly, *F_s_*(*q*,*t*) for tracer cells also displays two-step relaxation for the three values of *τ* investigated, as shown in Fig. 9b. Interestingly, the values of the first relaxation times are longer than for the corresponding dynamics associated with the cells. The results in Fig. 9b show that the dynamics of tracer cells is qualitatively similar to that calculated for the actual cells (see also Fig. 8a for comparison).

**FIG. 9.**
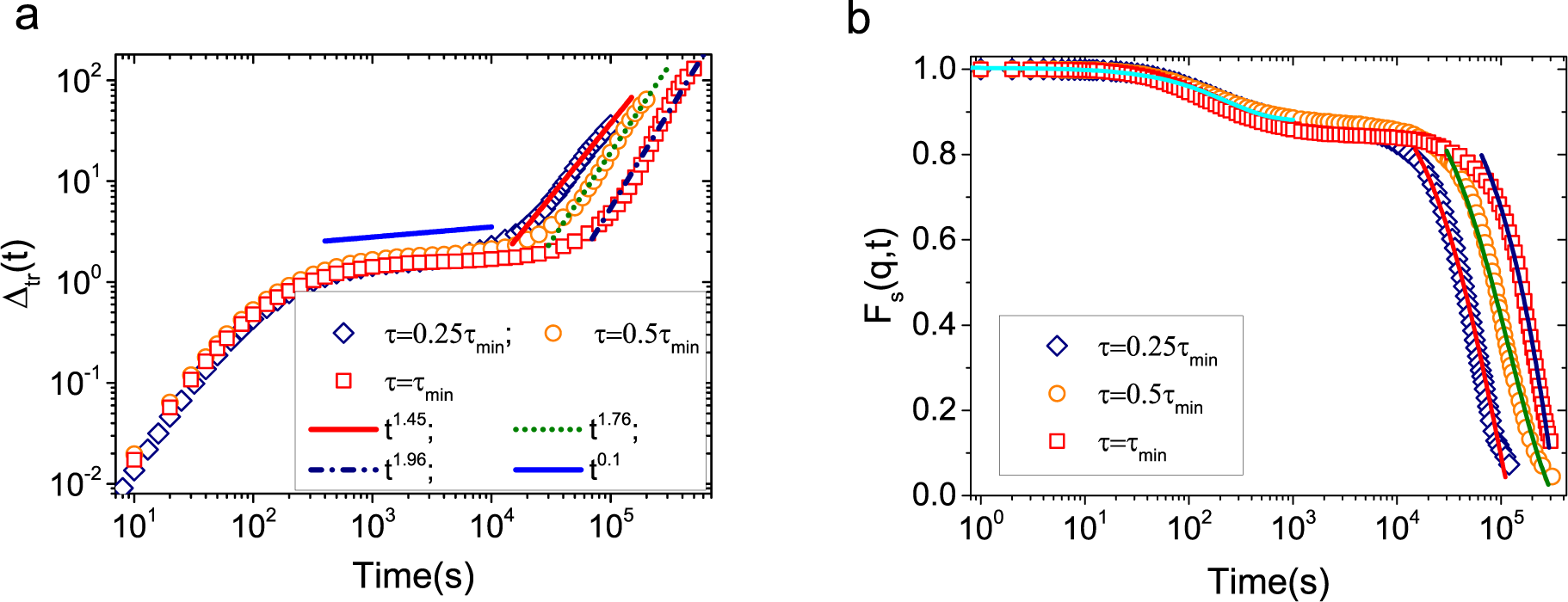
Dynamics of the tracers at different cell cycle times. **(a)**Time dependence of Δ(*t*) of tracer cells at different cell division times, *τ*. Fits to Δ(*t*) at intermediate time and long times are shown in the inset. **(b)** The self-intermediate scattering function, *F_s_*(*q,t*), for tracers. Biexponential fits to the decay of tracer *F_s_*(*q*,*t*) are shown by solid lines.

### Heterogeneity during tumor growth

The effect of glass or liquid-like state of tumor growth is illustrated by following the trajectories of individual cells in the growing tumor. Figs. 10a and fig. 10b highlight the trajectory of cells during a time of ≈ 3 days for the average cell division time of 15*τ_min_* and 0.25*τ_min_*, respectively. In the glass-like phase (intermediate times corresponding to *t*/*τ* < 1 for *τ* = 15*τ_min_*), the displacements are small, exhibiting caging behavior (Fig. 10a), resulting in the localization of the cells near their starting positions. On the other hand, cells move long distances and show sig-natures of persistent directed motion at the shorter cell cycle time in the long time superdiffusive regime corresponding to *t/τ >* 1 (see Fig. 10b). These observations suggest that the anisotropic growth of cells, manifested largely in the evolution of cells at the periphery of the tumor, depends on the cell growth rate, a factor that determines tumor virulency.

**FIG. 10.**
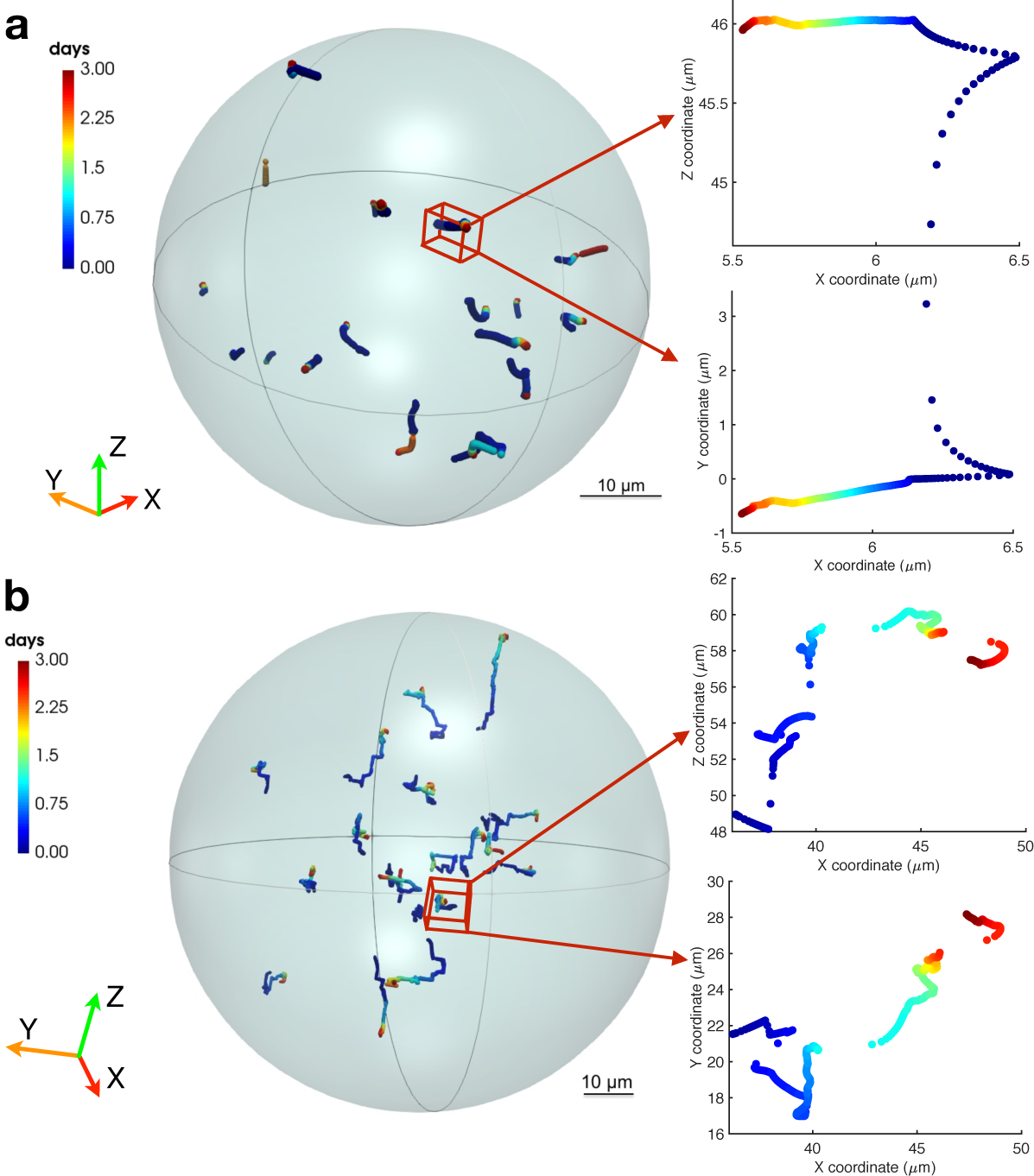
Trajectories displaying spatial heterogeneity. **(a)** Trajectories (randomly chosen from the whole tumor) for slowly growing cells are shown. Cell cycle time is 15Tmin. Dynamic arrest due to caging in the glass-like phase is vividly illustrated. **(b)** Trajectories for rapidly growing cells with cell cycle time *τ* = 0.25*τ_min_*. Displacements of the cells are shown over 3 days representing the initial stages of tumor growth in **(a)** and **(b)**. Two representative trajectories (time dependence of the *x – z* and *x – y* coordinates) for the labelled cells are shown on the right of **(a)** and **(b)**. Length in **(a)** and **(b)** is measured in units of *μm.* The two colored spheres in a and b show the approximate extent of the tumor.

To quantify spatial heterogeneity, we divided the tumor into two regions - interior and periphery. We note that such a division is not applicable in a system with periodic boundary conditions [27]. After obtaining the invasion distance Δ*r*(*t* = *t_E_*) (see Eq. (10)), we calculated the distance from the center of mass for the colony of all the initial cells that are alive at *t_E_*. Let us call this vector 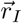. We grouped 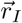 into two distinct categories: interior region if 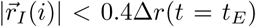 and boundary region if 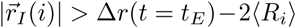. The average radius of all the cells in the tumor is denoted by ⟨*R_i_*⟩. We chose *t_E_* = 350,000s = 6.48*τ*. With this choice of *t_E_* we obtain good statistics allowing us to glean both the sub-diffusive and super diffusive behavior from the time dependence of Δ(*t*) (see Fig. 11). Once the initial cells are classified in this manner, we obtain their entire trajectory history and calculated their Δ(*t*). In Fig. 11, a plot of Δ(*t*) for the interior and boundary cells is shown. The dynamics associated with interior cells is sub-diffusive through their entire lifetime while the boundary cells show sub-diffusive motion at intermediate times and super-diffusive motion at long times. Interestingly, the cells at the boundary also show the intermediate glassy regime, which is *a priori* hard to predict.

**FIG. 11.**
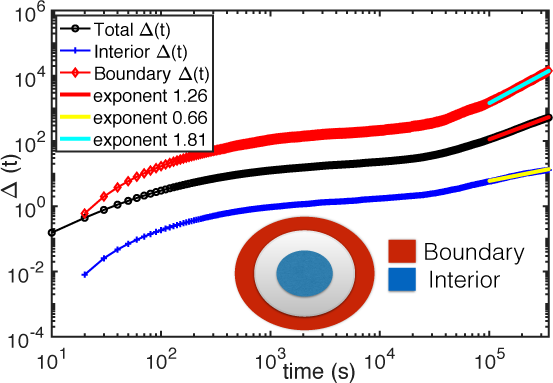
Plot of the mean squared displacement, Δ(*t*), for interior, boundary and the total initial cells. The interior cells exhibit sub-diffusive behavior through their entire lifetime. The cells at the boundary show both sub-diffusive motion at intermediate times followed by super-diffusive behavior at long-times. The interior Δ(*t*) is multiplied by 0.1 and the boundary Δ(*t*) by 10 for clarity. Cartoon depicting the way we have divided the tumor into interior and the boundary regions is also shown.

Because the nature of cell movement determines cancer progression and metastasis [58], it is critical to understand how various factors affecting collective cell migration emerge from individual cell movements (Figs. 10 & 12). Insights into cell migrations may be obtained by using analogies to spatial heterogenous dynamics in supercooled liquids [59, 60]. In simple fluids, the distribution of particle displacement is Gaussian while in supercooled liquids the displacements of a subset of particles deviate from the Gaussian distribution [59]. In Fig. 12a, the van Hove function of cell displacement (or the probability distribution of step size) is shown. The single time step distance covered by a cell is defined as |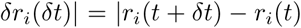|. By normalizing *δt* by the average cell cycle time i.e. *δr_i_*(*δt*/*τ* = 0.0074), we obtain a long-tailed *δr* probability distribution (*P*(*δr*)). The distribution *P*(*δr*), has a broad, power law tail cut off at large values of **δr*,* that depends on the cell cycle time. As we approach the glass-like phase for longer average cell cycle time, P(*δr*) is suppressed by an order of magnitude over a wide range of **δr*.* Interestingly, we do not observe an abrupt change in the behavior of P(*δr*) as the average cell cycle time is changed. The transition between glass-like and liquid-like regimes occurs continuously. To further analyze the displacement distribution, we fit the van-Hove function for squared displacements (*P*(*δr*^2^)) at normal cell division time (*τ_min_*) to both exponential and power law. The distribution is considerably broader than the Gaussian distribution (see Fig. 13 and Table II), providing one indication of heterogeneity [61].

**FIG. 12.**
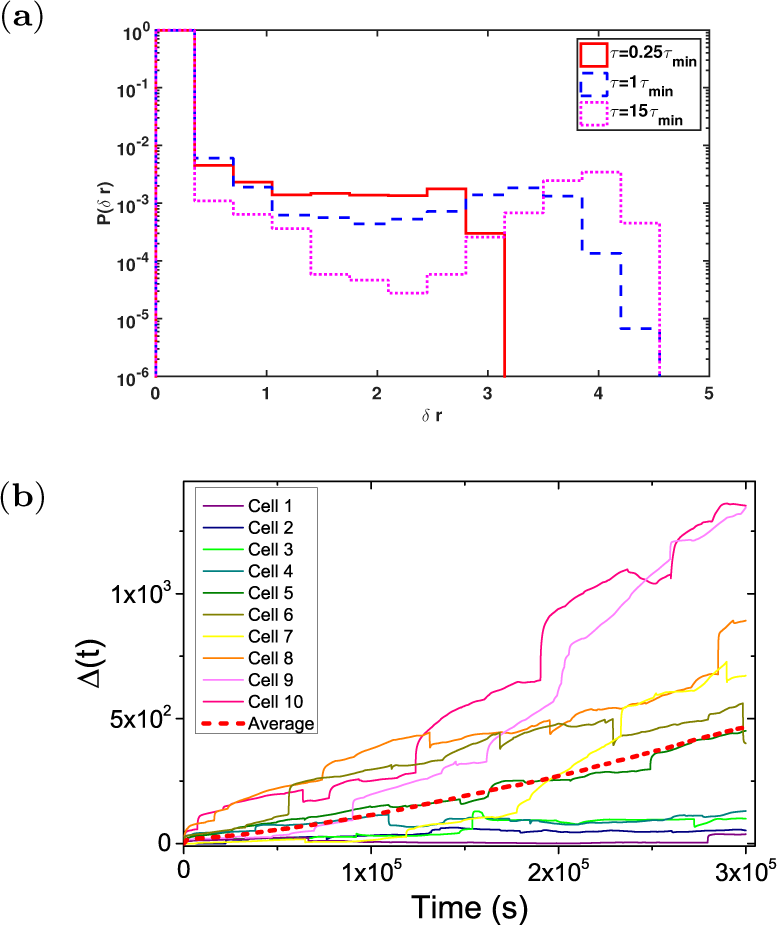
Quantifying spatial heterogeneity in tumor cell growth. (a) Probability distribution of distance *δr* (in unit of *μm),* moved by cells over *δt* = 100s, 400s, 6000s respectively for varying average cell cycle time *τ* = 0.25*τ_min_*, 1*τ_min_* and 15*τ_min_*. St is normalized by *τ* to 0.0074. (b) Time resolved squared displacements, Δ(*t*) (in unit of *μm^2^*), of individual cells in a model for growing tumor (*τ* = *τ_min_*). The average, shown as a dashed line for ≈ 800 such individual trajectories, is not meaningful because of dynamic heterogeneity.

**FIG. 13.**
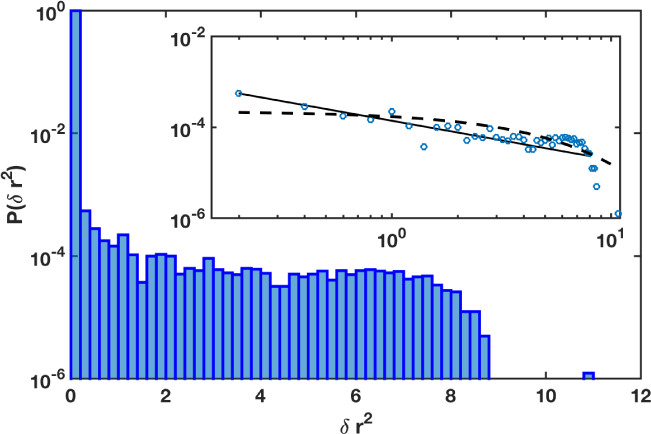
The probability distribution *P*(*δr*(*t*^2^_*i*_)) of cell displacements 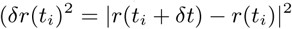 in units of *μm^2^*) at *δt* = 100s is shown. Cell trajectories until *t* = 5*τ* are analyzed at *τ* = 1*τ_min_*. The histogram was constructed by varying *t_i_* in *δr*(*t_i_*)^2^ for over 400 cell trajectories and after obtaining ≈ 10^6^ data points for *δr*(*t_i_*)^2^. For comparison, the inset shows fits to both exponential (dashed line) and power law (solid line). *P*(*δr*^2^) ~ *A* (*δr*^2^)^B^ fit the trend best where *A* is a constant and *B* is ~ 0.9. The striking non-Gaussian behavior, with fat power law tails, is one indication of heterogeneity. Goodness of fit can be assessed using the parameters listed in Table II.

**Table II:** The goodness of fit in the inset of Fig. 13. R-square value (middle column) and RMSE, the root mean squared error (last column) for power law and exponential fits are provided. The digits in parenthesis in the last column refer to powers of 10.

| Fit Type | $R^2$ | RMSE |
| --- | --- | --- |
| Power | 0.93 | 2.4(−05) |
| Exponential | 0.57 | 5.7(−05) |

Cell-to-cell phenotypic heterogeneity is considered to be one of the greatest challenges in cancer therapeutics [62, 63]. Within the context of our model, spatio-temporal heterogeneity in dynamics can be observed in tissues by analyzing the movement of individual cells. While the simulated time-dependent variations in the average mean-squared displacement is smooth, the movement of the individual cell is not (see Fig 12b). Cells move slowly and periodically undergo rapid ‘jumps’ or hops similar to the phenomenon in supercooled liquids [59, 60]. The squared displacement of individual cells (Fig 12b) vividly shows the heterogeneous behavior of different cells.

### Polarized tumor growth

From our simulations, we constructed a spatial map of the velocities of the individual cells in the tumor. Using these maps, we characterized the spatial heterogeneity in the dynamics in order to elucidate regions of coordinated activity in the movement of cells. Fig. 14a shows a snapshot of the spatial map of the single cell velocities. The velocity map, which spans more than eight orders of magnitude, reveals that there are cell-to-cell variations in the dynamics. More importantly, it also reveals the existence of spatial correlations between cell dynamics. In the tumor cross-section (Fig. 14b), faster moving cells are concentrated at the outer periphery of the tumor. By calculating the average magnitude of cell velocity as a function of radius, we show in Fig. 14c that faster moving cells are located at the outer periphery of the tumor quantitatively. We calculate the average velocity of cells at different radii of the tumor using,

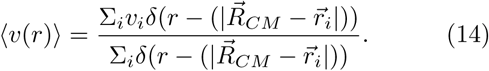

**FIG. 14.**
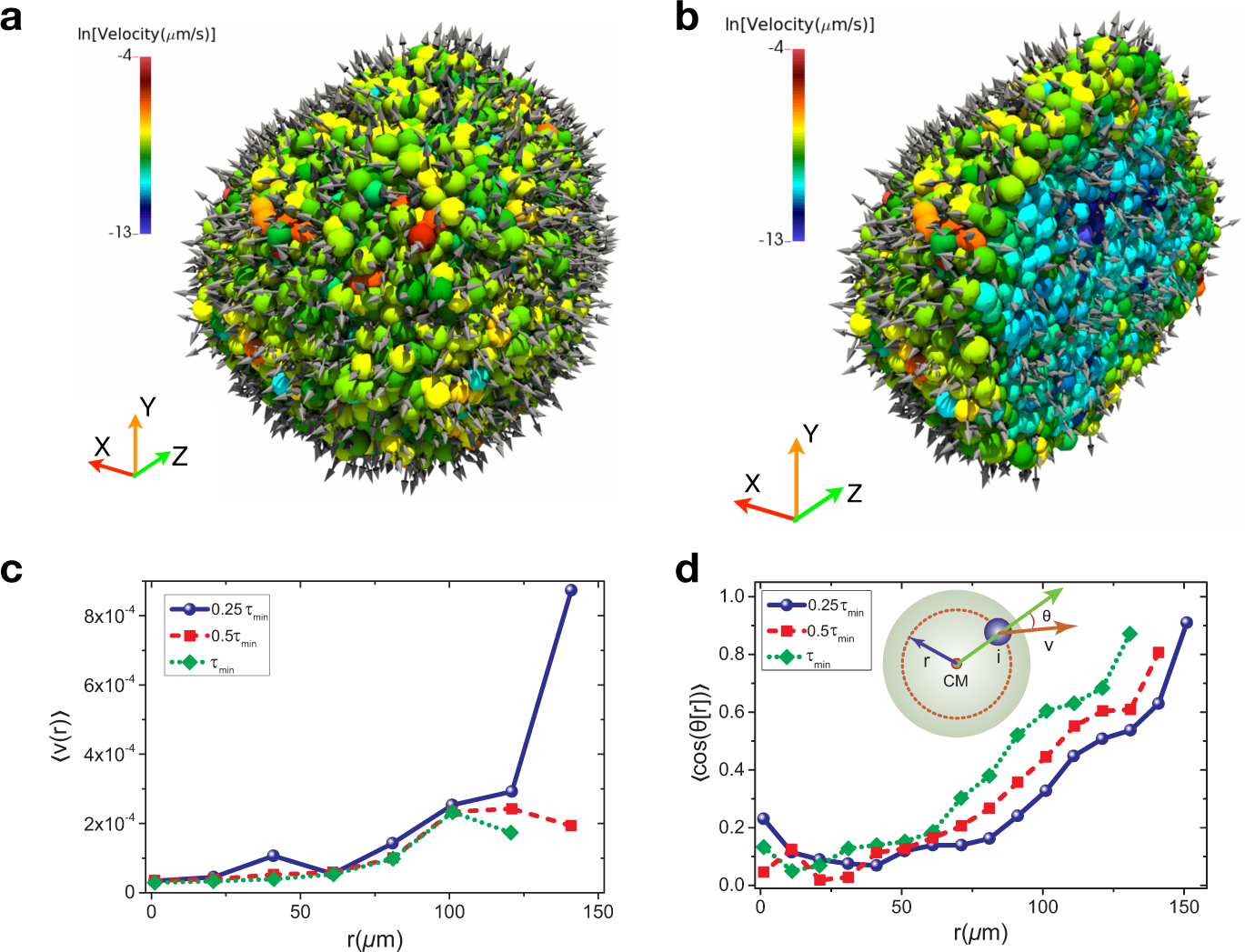
Heterogeneity in the tumor cell dynamics. **(a)** Instantaneous snapshot of a collection *N* ≈ 1.3 × 10^4^ cells at ≈ 3 days with *τ* = 0.25*τ_min_*. Colors indicate the different velocities of the individual cells (in *μ/s).* **(b)** Cross section through the clump of cells shown in Fig. 14a. Arrows denote the direction of velocity. **(c)** Average speed of the cells as a function of the tumor radius at different *τ*. Observation time is at 18.5*τ*, 14.8*τ* and 11.1*τ* for *τ* = 0.25*τ_min_*, 0.5*τ_min_* and 1*τ_min_* respectively. **(d)** Mean angle θ (see the inset figure) between cell velocity and a line through the center of the tumor to the periphery as function of the tumor radius at different *τ*. Observation time is the same as in **c**.

Mean angle *θ* between cell velocity and the position vector with respect to the center of the tumor plotted in Fig. 14d further illustrates that cell movement becomes persistently directed outward for cells closer to the outer layer of the tumor. To calculate the radius-dependent average polarization in cell velocity, we first define a vector pointing from the center of mass of the tumor to the cell position 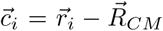 (see the green arrow in the inset of Fig. 14d). The angle *θ* (see the inset of Fig. 14d) between each cell velocity (orange arrow) and the vector (green arrow) from the center of mass to the tumor periphery can be calculated from 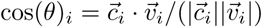. The average of this angle as a function of radius is calculated using,

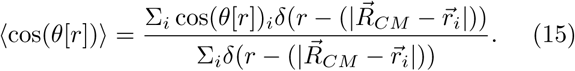

Arrows indicating the velocity direction show that cells in the periphery tend to move farther away from the center of the tumor as opposed to cells closer to the center of the tumor whose direction of motion is essentially isotropic. This prediction agrees well with the experiments [33], which showed that cells at the periphery of the tumor spheroid move persistently along the radial direction, resulting in polarized tumor growth.

The results are presented in Fig. 14d in the main text. The distribution of the *θ* angle at different distances (*r*) (see Fig. 15a) also illustrates that cell movement is isotropic close to the tumor center, while they move outward in a directed fashion (see the peak of the histogram in blue) at the periphery of the tumor. To quantify the heterogeneity in cell velocity, we plot the probability distribution of the velocity magnitude (normalized by the mean velocity - ⟨*v*⟩), *P*(|*v*|/⟨*v*⟩), (Fig. 15b) accessible in experiments using direct imaging or particle image velocimetry methods [51, 52]. There is a marked change in the velocity distribution as a function of cell cycle time. At the longer cell cycle time (*τ* = 15*τ_min_*), corresponding to the glass-like phase, P(|*v*|/⟨*v*⟩) distribution is clustered around smaller values of |*v*|/⟨*v*⟩ while quickly decaying to zero for higher velocities. For the shorter cell cycle time, the velocity distribution is considerably broader.

**FIG. 15.**
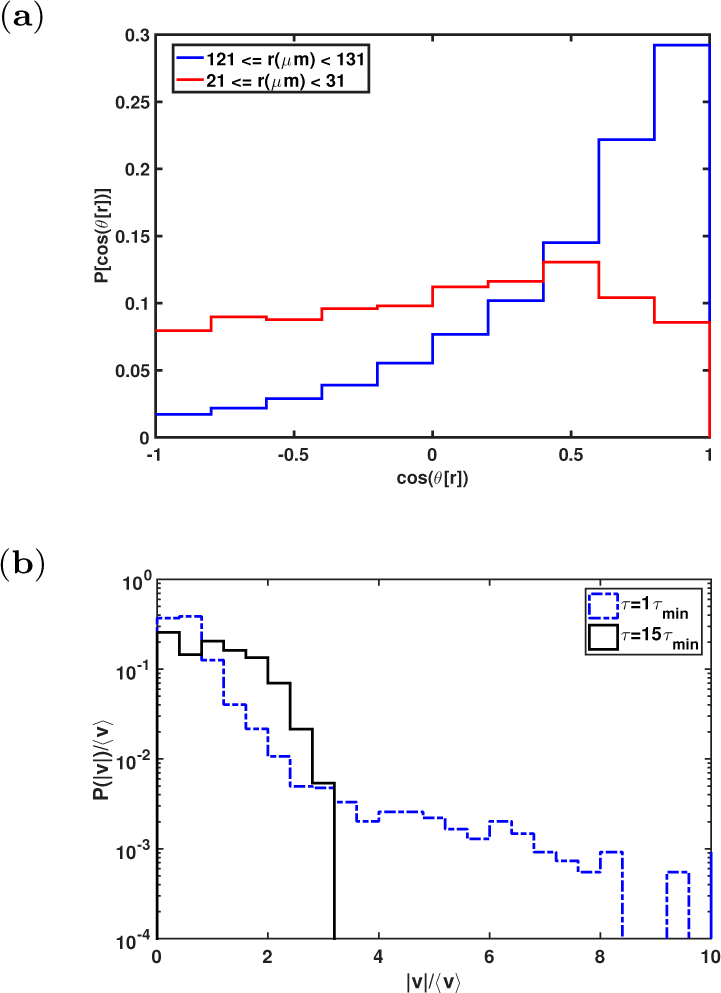
Heterogeneity in the tumor cell dynamics(continued). **(a)** Distribution of the angle (θ) at different distances (*r*) from tumor center at ≈ 3 days (*τ* = 0.25*τ_min_*). **(b)** Probability distribution of the cell speed normalized by mean cell velocity, ⟨*v*⟩, at two different cell cycle times at the long time regime (*t* = 5*τ*).

The broader velocity distribution indicates the presence of more invasive cells within the tumors characterized by high proliferation capacity.

### Consistency with experiments

We show here that the minimal model captures the three critical aspects of a recent single-cell resolution experiment probing the invasion of cancer cells into a collagen matrix [33]: (*i*) Ensemble-averaged mean square displacement of individual cells exhibit a power-law behavior at long times (Δ(*t*) ~ *t^α^* with *α* ≈ 1.40 ± 0.04 from experiments compared with simulation results in Fig. 5(a) with *α* ≈ 1.26±0.05) indicating that, on an average, directed rather than random cell motion is observed. (*ii*) Cells exhibit a distinct topological motility profiles. At the spheroid periphery cell movement is persistently along the radial direction while stochastic movement is observed for cells closer to the center. Such spatial topological heterogeneity is well-described as arising in our model from pressure dependent inhibition (see Figs. 10, 12 & 14). (*iii*) The highly invasive spheroid boundary (deviating from what would be expected due to an isotropic random walk) as experimentally observed is qualitatively consistent with simulation results (see Fig. 6).

A salient feature of the dynamics of living cells is that birth and death processes break number conservation, having consequences on their collective behavior [64]. To account for these processes leading to the super-diffusive behavior at long times, we establish a field theory based on stochastic quantization that account for the physical interactions of the cells as well as birth and death processes. Simulations and theory suggest a mechanism of the plausible universality in the onset of super-diffusive behavior in tumor growth and unrelated systems. Remarkably, the theory predicts the dynamics of invasion at all times that are in good agreement with recent experiments [33].

### Onset of super-diffusion depends on cell division time

In previous studies [26, 27], fluidization of tissues due to cell division and apoptosis was observed at the homeostatic state. Our work shows that a glass-to-fluid transition is driven by cell division at non-steady states and under free boundary conditions, relevant during early stages of cancer invasion. The transition from glass to fluid-like behavior is determined by the average cell division time. Super-diffusion of cells in the mean-squared displacement due to highly polarized tumor growth is observed on a time scale corresponding to the cell division time with a universal scaling exponent *α* = 1.26 ± 0.05.

### Comparison with previous studies

The startlingly contrasting results that we find for the dynamics of tracer cells and live cells compared to the results reported elsewhere [27] could arise for the following reasons: (i) In our simulations, we use free boundary conditions and the tumor grows continuously (with birth rate being always higher than the death rate). The effect of a free boundary is particularly pronounced at the periphery of the tumor where the cells undergo rapid division. (ii) Imposing a birth rate that depends on the local cell density (Ref. [26]) or on the number of nearest neighbors as in Ref. [27] eventually results in a homeostatic state where birth and death are balanced. The super-diffusive behavior, observed in our study would not be present, when there is a possibility of reaching a homeostatic dense liquid-like state. In our model, this situation could be mimicked by arbitrarily increasing the cell cycle time. For example, when cell cycle times are very long *τ* > ~ 10 *τ_min_*, the super-diffusive MSD exponent in the long time regime begins to deviate from ≈ 1.3 to lower values (see Fig. 5(a)). (iii) Even in our model, simple diffusion at long times is obtained if the death rate is modified. In Fig. 16, normal diffusion (red symbols) is observed at long times when the death rate is modified to

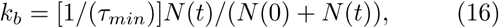

where *N*(*t*) and *N*(0) are the tumor size at time *t* and *t* = 0 respectively. At *t* = 0, cells have a higher birth rate, *k_a_* = 1/(*τ_min_*), compared with the death rate (*k_b_/k_a_* =0.5).

**FIG. 16.**
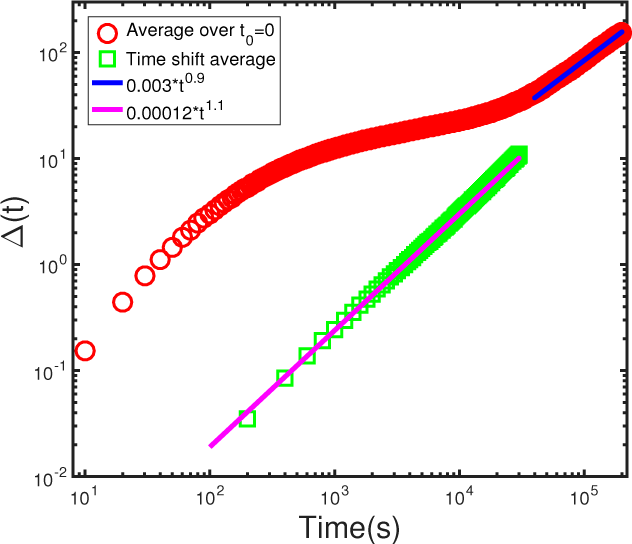
The mean-squared displacement (Δ(*t*)) of cells when cell death rate depends on time. The red circles shows the results obtained by averaging over initial cell position, Δ(*t*) =< [*r*(*t*)–*r*(*t*_0_ = 0)]^2^ >. The green squares show the results calculated using time shift average, Δ(*t*) =< [*r*(*t_i_* + *t*) – *r*(*t_i_*)]^2^ >. The solid lines show power-law fits to the simulation data. Normal diffusion results because the use of Eq.16 leads to homeostasis at long times.

As the tumor grows, the death rate becomes higher and a homeostatic state is reached once the birth and death rates are balanced, giving rise to normal diffusion as found in Ref. [26, 27]. Therefore, the super diffusive behavior can only be found if birth and death processes are not balanced, the regime which is the focus of our study. Most importantly, if there is a mechanism for reaching homeostasis by balancing birth and death rates or making the cell division time arbitrarily long we predict that normal diffusion would result, as shown here using Eq.16 and previously found elsewhere [26, 27].

### Lack of time translational invariance

The field theory shows that the dynamics is not time translationally invariant, which is supported by simulation of Δ(*t*). Fig. 17 shows two different methods used to calculate the Δ(*t*): (i) the definition of MSD,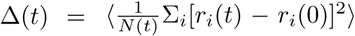, which always utilizes the original position of cells (at the initial simulation time) (*r*(0)). Here, *N*(*t*) is the number of initial cells and ⟨…⟩ is the average over multiple simulation runs; (ii)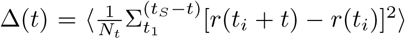, a time shift average (varying *t_i_*), used routinely in simulations of periodic systems with *N_t_* being the number of possible time intervals for a given *t*. Here, ⟨…⟩ denotes average over initial cells. In generating the results in Fig. 17 we chose *t_S_* = 500,000s.

**FIG. 17.**
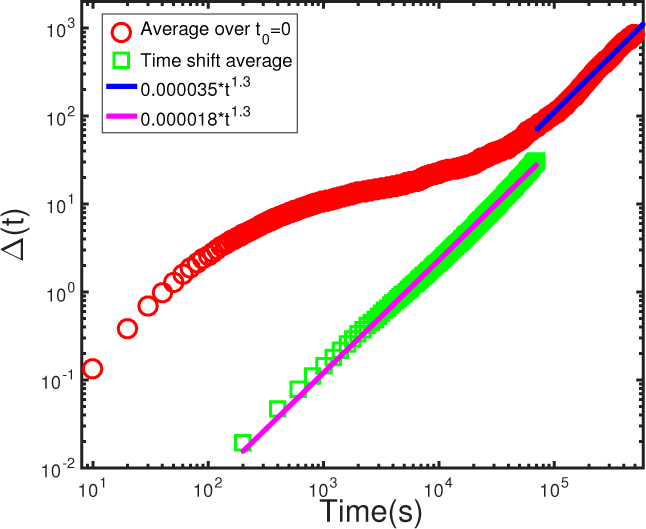
Mean squared displacement, Δ(*t*), of cells as a function of time. The red circles show the results obtained by averaging over the initial cell positions, Δ(*t*) = ⟨[*r*(*t*) – *r*(0)]^2^⟩. The green squares are the results under time shift average, Δ(*t*) = ⟩[*r*(*t_i_* + *t*) – *r*(*t_i_*)]^2^⟩. The solid lines show power-law fitting of the simulation data. The long time super-diffusive behavior is evident in both the plots.

These two methods for computing Δ(*t*) produced different results, as shown in Fig. 17. The inequivalence of the two methods in obtaining Δ(*t*) shows that the system with free boundary conditions violates time translational invariance. The intermediate regime with sub-diffusive behavior (red circles) using method (i) disappears when the second method (ii) is used to compute the Δ(*t*) (green squares). It is also the reason why we focused on initial cells for Δ(*t*) calculations. In calculating Δ(*t*) using the method (ii) commonly employed in simulations of periodic systems, large amount of statistics is extracted for the super-diffusive regime (as it has larger time range on the order of ≈ 10^5^s). Therefore, when we average Δ(*t*) over the various time intervals, the beginning time regime (which is comparatively short ≈ 10^4^s) is suppressed. It should be noted, however, that the time shift averaged method of computing Δ(*t*) also clearly shows evidence for super diffusive behavior over three decades in time (Fig. 17).

## CONCLUSIONS

Heterogeneity is a hallmark of cancer [65]. It is difficult to capture this characteristic of cancers in well-mixed models that exclude spatial information. An important signature of cell dynamic heterogeneity - large variations in the squared displacement of cells in the tumor - is observed in our simulations. We find a broad velocity distribution among tumor cells driven by cell growth rate. The formation of spatial niches, with tumor periphery and center as being topologically distinct is characterized by differences in proliferative and cell signaling activities. Such a distinct behavior, alluded to as the driving factor behind intratumor heterogeneity (ITH) [66, 67], is not well understood. Our results predict that the pressure dependent inhibition of cell growth to be the critical factor behind the development of distinct topological niches, implying that the dynamics of cells is dependent on the microenvironment [66, 67]. Cells closer to the center of the tumor spheroid are surrounded by many other cells causing them to be predominantly in the dormant state and can move in random directions, while cells closer to the periphery can divide and move in a directed manner by pushing against the extracellular matrix, thus promoting tumor growth and invasiveness. We provide experimentally testable hypotheses on the signatures of heterogeneity - the onset of ITH could occur at very early stages of tumor growth (at the level of around 10,000 cells).

Although the context of our work being rooted in understanding tumor growth, we expect our model to be relevant to the study of soft glassy materials. The motion of cells in our model is surprisingly consistent with the complex motion of bubbles in a foam, also shown to be super-diffusive [36] with the MSD exponent of *α* ≈ 1.37 ± 0.03, consistent with both our theoretical predictions (*α* ≈ 1.33) and simulation results (*α* = 1.26±0.05). The bubbles are characterized by birth and death processes and pressure dependent growth, which we predict to be the driving factors behind super-diffusive behaviors observed in these diverse systems. Emergence of underlying similarities in the motion of constituent particles between living systems, such as cells, and soft glassy materials, such as foams, suggest that many of the shared, but, as of yet unexplained dynamic behavior may emerge from a common underlying theme - an imbalance in the birth and death processes and pressure dependent growth inhibition.

## ACKNOWLEDGMENTS

We acknowledge Anne D. Bowen at the Visualization Laboratory (Vislab), Texas Advanced Computing Center for help with figure and movie visualizations. We are grateful to Mauro Mugnai, Naoto Hori and Upayan Baul for discussions and comments on the manuscript. This work is supported by the National Science Foundation (PHY 17-08128 and 16-32756). Additional support was provided by the Collie-Welch Chair (F-0019). A. N. M and X. L. contributed equally to this work.

## Appendix A: Simulation Test

### Effects of random forces

The neglect of random forces, which should be taken into account to satisfy the FDT, might seem like a drastic simplification. There are, however, two considerations. First, the tumor growth model involves birth and apoptosis. Hence, it behaves like an active system. Indeed, the theory outlined in Section C shows that under these conditions FDT is not satisfied, forcing us to adopt the stochastic quantization methods to compute response and correlation functions (Fig. (19)). Second, from practical considerations we note that the cellular diffusion constant is 10^−4^ *μm^2^*/s or smaller [29], resulting in only small displacements for a large fraction of cells.

**FIG. 18.**
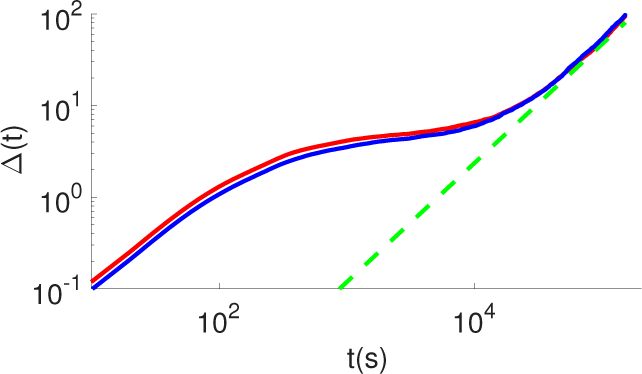
Mean square displacement, Δ(*t*), with (blue) and without (red) random noise. The slope obtained from the long time limit are both 1.3 (dashed green). The two curves are almost identical, thus justifying the neglect of the random noise (second term in Eq. (A1)) in the simulations.

**FIG. 19.**
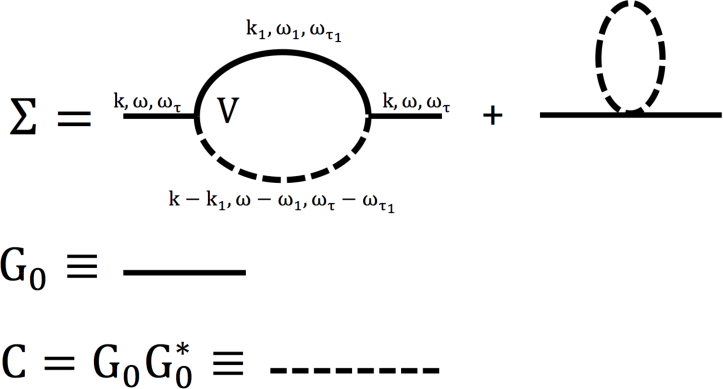
The diagrams correspond to perturbation expansions of the theory (Eq. (C3)) in which the dynamical equations for the density field is expressed in fictitious time. Self-energy term (Σ) is obtained by contracting the two density *ρ* fields. The first diagram is the two loop contribution generated from the first order term (contains two *ρ* fields) in the time-dependent equation for the density fields. The second diagram, with one loop contribution from the second order term (contains three *ρ* fields), resulting in the correction to *ω*^2^ + {*C*_0_*k*^2^*U*(**k**) – (*k_a_* – 2*k_b_C*_0_)}^2^ does not have any new momentum dependance. Hence, only the first term is significant in producing the scaling results.

In order to verify that the contributions to the dynamics arising from the random noise is small, we modified Eq. (7) to include the random forces,

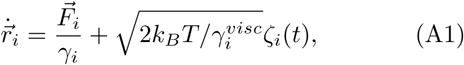

where *k_B_* is the Boltzmann constant, *T* the temperature and ζ is white noise with zero mean and variance, 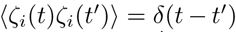. The corresponding diffusion constant, *k_B_T*/*γ*^*visc*^, is small. Thus, inclusion of random force has no consequence on the dynamics of tumor evolution. The results for Δ(*t*) as a function of *t* obtained using Eqs. (7) and (A1) are identical (Fig. 18).

## Appendix B: MOVIES

In order to visualize the dynamic growth of the tumor we generated movies from the simulations. They demonstrate vividly the polarized growth of the tumor, which we have quantified using various measures in the main text.

### Supplementary Movie 1. 3D growth of tumor

This movie shows the three dimensional growth of the tumor over ≈ 8 days. Each frame is at 1000 seconds. The cell cycle time *τ* = *τ_min_*. Colormap indicates the life time of the cells. Newborn cells are shown in blue and older cells that have lived longer are in red (color bar in video shows cell lifetime in seconds). Cell division and death events are explicitly depicted.

### Supplementary Movie 2. Cross section view through the growing tumor spheroid

Illustration of an alternate view of the growing tumor shown in Supplementary Movie 1. Cells with longer lifetimes are mostly localized near the center of the tumor with some of them moving to the periphery. Newly born cells are mostly located in the periphery and division events are amplified in the periphery compared to the center of the tumor. Color bar shows cell lifetime.

### Supplementary Movie 3. Moving clip through a tumor showing velocity heterogeneity

This video visualizes the velocity heterogeneity within the tumor. Colormap indicates the speed of cells (shown in log scale) and the direction of velocity is indicated by an arrow. The video begins with a snapshot of the tumor after ≈ 3 days of growth at *τ* = 0.25*τ_min_*. A clip moving through the tumor shows the velocity distribution of cells over different slices. It is clear that cells move slowly closer to the center while faster moving cells are mostly in the periphery. Direction of the velocity is more randomly oriented in the tumor center but is mostly polarized outward as the periphery is approached.

## Appendix C: Theory

The behavior of the Mean Square Displacement (Δ(*t*)), especially the time dependence of Δ(*t*) at intermediate and long times, can be theoretically obtained for the tumor growth model mimicking the one used in the simulations. We consider the dynamics of a colony of cells in a dissipative environment with negligible inertial effects. The interaction between cells is governed by adhesion and excluded volume repulsion. The equation of motion for a single cell in a dissipative environment with negligible inertial effects. The interaction between cells is governed by adhesion and excluded volume repulsion. The equation of motion for a single cell *i* is,

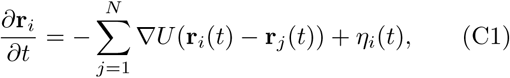

where *U* contains the following form of repulsive interactions with range λ, and favorable attractive interactions between cells with range *σ*,

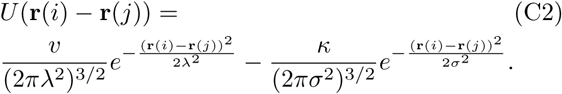

*v* and *k* above are the strengths of the repulsive and attractive interactions, respectively. The *κ* parameter in Eq. C2 mimics adhesion between cells. The noise (*η_i_* in Eq. (C1)) is uncorrelated in time.

The simplified form for *U*, which captures minimally the interactions between cells but differs from the more elaborate model used in the simulations, allows us to obtain analytical results for Δ(*t*) as a function of t. In terms of the density field of a cell, 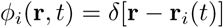, a closed form Langevin equation for the density, 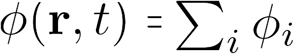 can be obtained using the approach introduced by Dean [68]. In order to study tumor cell dynamics, we extend the model phenomenologically to describe both cell division and death, and introduce a noise term that breaks the cell number conservation. These crucial features needed to describe tumor growth can be investigated using the Doi-Peliti (DP) formalism [69, 70], introduced in the context of reaction-diffusion processes. A related approach was used recently by Gelimson and Golestanian [71] to describe collective dynamics in a dividing colony of chemotactic cells.

We use a scheme to study the interplay between stochastic growth and apoptotic process, and use it to derive a Langevin type equation for logistic growth. The birth reaction, 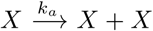, occurs with the rate constant *k_a_* for each cell, and the backward reaction (apop-tosis) 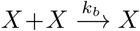 occurs with rate *k_b_*. By incorporating birth and apoptosis, and assuming that the density fluctuates around a constant value, *ϕ*(***r**, t*) = C_0_ + *ρ*(**r**, *t*), we obtain the approximate equation for the density fluctuation, which in Fourier space reads,

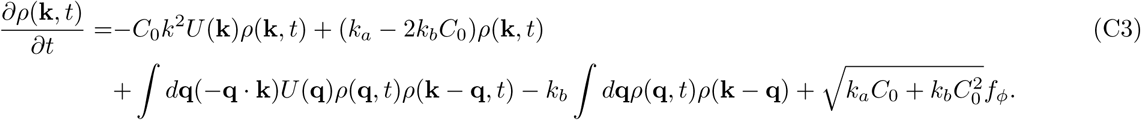

We derived Eq. (C3) by expanding the density to lowest order in 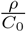 non-linearity. The noise *f_ϕ_* satisfies 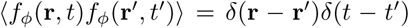. In the hydrodynamic, *k →* 0 and *t* → ∞ limit, the first and third terms in the RHS of Eq. (C3) vanish, and hence the scaling behavior of Δ(*t*) at long times is determined solely by the death-birth terms.

The scaling of Δ(*t*) can be obtained by treating the non-linear terms in Eq. (C3) perturbatively using the Parisi-Wu stochastic quantization scheme [55, 72, 73], which is needed because Fluctuation Dissipation Theorem (FDT) is not satisfied in Eq. (C3) due to cell birth and death processes. In order to outline the essence of the theory, let us consider the probability distribution corresponding to the noise term given by

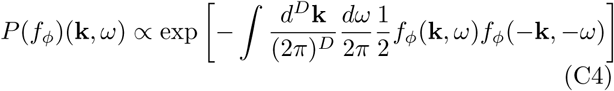

By re-expressing *P*(*f_φ_* (**k**,*ω*)) in terms of P(*ρ*(**k**,*ω*)), a Langevin equation of motion in the fictitious time, *τf*, may be derived in which FDT is satisfied. Consequently, in the τf → ∞ limit, the distribution function P(*ρ*(**k**,*ω*)) ∝ exp(–S(**k**,*ω*)), where an expression for the effective action *S*(**k**,*ω*) is derivable from Eqs. (C3) and (C4). Using this formalism, the Green’s function can be obtained using perturbation theory by solving the Dyson equation,

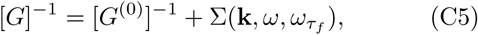

where *ω_τf_* is the frequency related to *τ_f_* and 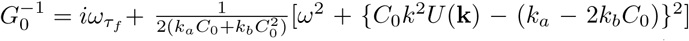. Diagrammatic representation of self energy term 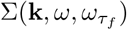 is shown in Fig. 19 to one loop order. We obtain,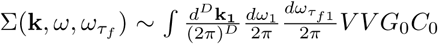, where the vertex term is of the form: 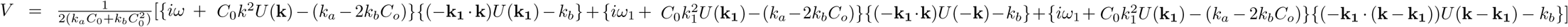, the correlation 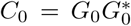, and *D* is the spatial dimension. After computing the self energy to second order in non-linearity, Eq. (C5) can be written as,

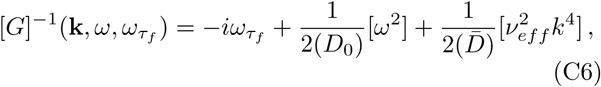

where 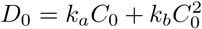. The above equation allows us to determine an effective coefficient 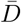 from *G*^−1^ (**k**, 0,0),

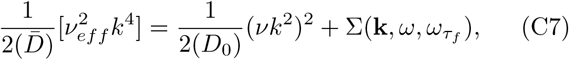

with *v* = *C*_0_*U*(**k**). In obtaining Eq. (C7), needed for calculating the scaling of Δ(*t*) in the intermediate time, the strength of the interactions are such that *C*_0_*k*^2^*U*(**k**) dominates over (*k_a_* – 2*k_b_C*_0_). Expanding *v_eff_* about *v* and 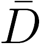 around *D*_0_, and noting that the renormalization of *v* dominates, we write using Δ*v* = *v_eff_* – *v*,

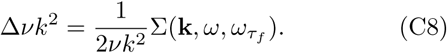

In the spirit of self-consistent mode coupling theory, we now replace *v* by Δ*v* in the self energy term Σ(**k**, *ω, ω_τf_*), use *G* as given by Eq. (C6), and the correlation function *C* = *GG*\*, as follows from the FDT. According to scale transformation, *ω ~ k^z^*, *ω_τf_ ~ k^2z^*, *G ~ k ^−2z^*, *C ~ k^−4z^* and the vertex factor *V* ~ *k*^*z*+2^. The self energy term in Fig. (19) can be written as 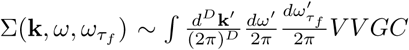 (Fig. (19) provides a diagrammatic representation of the theory). By carrying out the momentum count of Σ(*k,ω,ω_τf_*), and noting that *vk^2^ ~ k^z^*, we find Σ(*k,ω,ω_τf_*) ~ *k*^*D*−*z*+4^. Using Eq. (C8), we obtain *k^z^*+^2^ ~ *k*^*D*−z+4^, leading to 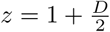.

The scaling of Δ(*t*) at intermediate and long times may be gleaned using the relation *C* = (1/*ω_f_*)Im *G*. Assuming dynamic scaling holds, the single cell mean-square displacement should behave as,

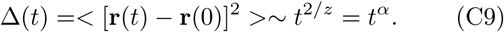

In 3D, 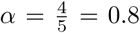, implying Δ(*t*) should display sub-diffusive behavior. The theoretical prediction is in accord with the behavior of Δ(*t*) in the caging regime. In the long time limit, the non-linearity due to death-birth dominates over mechanical interactions (∝ *U*(*k*)). A similar procedure, as mentioned above, produces the dynamic exponent *z* = *D*/2. In this regime, *α* = 1.33, implying super-diffusive motion, a prediction that is also in agreement with our simulations and experimental results [33]. Thus, the theory explains the simulation results, and by extension the experimental data, nearly quantitatively.

